# A MUC5B polymorphism associated with Idiopathic Pulmonary Fibrosis mediates overexpression through decreased CpG methylation and C/EBPβ transcriptional activation

**DOI:** 10.1101/2019.12.18.880674

**Authors:** Amaranta U. Armesto-Jimenez, Ari J. Arason, Olafur A. Stefansson, Gunnar Gudmundsson, Thorarinn Gudjonsson, Magnus K. Magnusson

## Abstract

Idiopathic pulmonary fibrosis is a progressive and fatal lung disease of unknown aetiology. The strongest genetic risk factor associated with IPF development is a *MUC5B* promoter polymorphism (*rs35705950*). However, the mechanism underlying its effects remains unknown. In this study we have focused on the molecular consequences of the polymorphism on the regulation of MUC5B expression. We have identified a combined mechanism involving both methylation and direct transcriptional regulation mediated by the polymorphic variant on MUC5B overexpression. Our results demonstrate that the minor allele (T) associated with *rs35705950* disturbs a DNA methylation site, directly increasing MUC5B expression. Furthermore, this same variant also creates a novel binding site for the transcription factor C/EBPβ leading to transcriptional activation of *MUC5B*. Our findings provide a novel insight into the regulatory effects of the IPF risk allele, *rs35705950* and identifies C/EBPβ as an important regulatory factor in the development of IPF.

## Introduction

Idiopathic pulmonary fibrosis (IPF) is an irreversible interstitial lung disease characterized by a progressive scarring of lung parenchyma often leading to a fatally declining lung function. IPF incidence has been estimated to be around 75/100,000, affecting 5 million people world-wide [1–3]. Furthermore, IPF is believed to be an under diagnosed disease [4–6].

Due to the complexity and progressive nature of the disease, available treatments are limited and have only a modest impact on IPF progression. Until recently, only lung transplant has been proved to increase survival [1]. In recent years, improvement in development of new therapies has met some success, as two novel drugs have been introduced, pirfenidone and nintedanib. Both drugs are believed to target profibrotic signalling pathways in IPF. Pirfenidone inhibits TGF-β1 production possibly by inhibiting the upregulation of HSP47 and Col1 RNA in fibroblasts [7]. Nintedanib is a potent small-molecule receptor tyrosine kinase inhibitor targeting platelet-derived growth factor receptors *(PDGF-R)*, fibroblast growth factor receptors *(FGFR)* and vascular endothelial growth factor *(VEGF)*-family tyrosine kinase receptors [8].

Both genetic and environmental factors are believed to contribute to the onset and progression of the disease. Several environmental factors have been associated with IPF, including exposure to metal and wood dust [9–12], viruses [13–15], drugs [16–18], and cigarette smoke [19–21]. Recent findings point to genetic factors as major triggers of IPF. Rare mutations found in 6 genes (*TERT* [22, 23], *TERC* [22, 23], *RTEL1* [24, 25], *PARN* [25], *STPC* [26, 27] and *SFTPA2* [28]), with associated variants in 11 different loci [29] indicate that the telomerase pathway and surfactant protein genes play a key role in IPF pathogenesis. However, these mutations only explain a small proportion of IPF cases.

Two large genome-wide association studies (GWAS) have been conducted on pulmonary fibrosis (familial and sporadic) [30, 31]. Both studies showed the most important genetic risk to be conferred by a common G-to-T risk variant in the upstream region of *MUC5B* (*rs35705950*). The frequency of the risk allele (T) is ~35% among European ancestry cases, compared with ~9% of European ancestry controls [32, 33]. These studies furthermore identified other common variants, including three polymorphisms in the *TOLLIP* gene and a desmoplakin (*DSP*) intron variant [31]. These studies concluded that the genome-wide variants account for 30-35% of the IPF risk, suggesting an important role for these common genetic variations in the disease aetiology [33].

Deciphering the molecular and cellular effects of non-coding SNP’s has proven to be notoriously difficult. With the thousands of common SNPs associated with many common diseases only a few have led to a clear molecular understanding [34–36]. There are many possible explanations for this. Most often there is a complex network of SNPs in varying degrees of linkage disequilibrium all associated with the phenotype or disease [36]. Resolving the true informative SNP within this complex network of variants can be challenging. Another explanation for the difficulty is connecting a non-coding variant to a target gene or regulatory region mediating the effects of the genetic association. Even though the closest gene may be a good candidate, the effects can be on genes further away. This can sometimes be resolved through expression-quantitative trait loci (eQTL), i.e. if the sequence variant influences the expression level of one or more genes.

The *MUC5B* promoter variant *rs35705950* is the strongest known risk factor (genetic and otherwise) for the development of IPF [32, 33]. This variant is unusual as it is relatively simple, with no other variants in strong linkage disequilibrium and carries a strong eQTL association with MUC5B expression in lung. The odds ratio (OR) associated with carrying one allele of the variant is 4.5-6.6, while homozygosity leads to an OR of 9.6-20.2 [30, 31, 37–41]. Furthermore, carriers of the *MUC5B* variant are also at risk of getting subclinical interstitial lung disease based on screening with high-resolution CT [4]. Interestingly, recent studies have examined the relevance of *MUC5B* variation in other ethnic groups, showing that the frequency of *rs35705950* minor allele (a G-to-T SNP) is 11, 8 and 1% among European, South Asian and East Asian populations, respectively [38, 42, 43], and it is almost non-existent in Africans [44]. Despite the different prevalence, the evidence indicates that the risk of developing IPF associated with the variant is comparable to the risk observed in European population [4, 31].

MUC5B (Mucin 5B) is a highly glycosylated protein, expressed throughout the upper and lower respiratory tract. In human upper airways MUC5B is predominantly expressed in nasal and oral gland secretions, while in tracheobronchial conducting airways it is expressed in surface epithelium and submucosal glands, where it is less predominant than the related MUC5AC in surface epithelia. In bronchioles MUC5B predominates, while MUC5AC is less expressed. Under disease conditions, the MUC5B expression pattern changes dramatically, it becomes overexpressed, showing significantly higher expression than MUC5AC [1, 45, 46]. The pattern suggests an important role for MUC5B as a host defence barrier. The G-to-T *rs35705950* variant is in a presumed regulatory region 3 kb upstream of the transcription start site in the *MUC5B* gene on chromosome *11p15*. A DNase I hypersensitivity site overlaps the variant location suggesting an important regulatory role. Furthermore, the *MUC5B* variant lies within a cluster of CpG sites and CpG DNA methylation has been shown to affect *MUC5B* expression [47].

Even though the *MUC5B* variant has a strong and consistent genetic association with IPF and the variant seems to positively regulate MUC5B expression, the mechanism underlying its role in IPF is poorly understood. It has been hypothesized that, due to the large size of the MUC5B protein, its production may carry a significant metabolic stress, which can interfere with differentiation of airway stem cells [33]. It should also be emphasized that due to the increased expression level there could also be secondary problems with post-translational modifications, such as glycosylation. In a mouse model of intestinal inflammation aberrant mucin assembly has been shown to cause ER-stress through activation of the unfolded protein response leading to inflammation, apoptosis, and wound repair [48]. ER-stress has been shown to be involved in IPF, specifically in cases caused by mutations in surfactant proteins, another protein that is highly glycosylated [49]. Other proposed mechanisms consider the possibility that IPF is a mucociliary disease caused by recurrent injury/inflammation/repair at the bronchoalveolar junction, as MUC5B overexpression might cause a reduction in mucociliary function and retention of particles leading to lung injury [33].

In this study, we aim to define the molecular mechanisms involved in mediating the effects of the *MUC5B* variant (T allele) on *MUC5B* expression. We use two immortalized human bronchial epithelial cell lines (BCi_NS1.1 and VA-10) and the well-established adenocarcinoma cell line A549 to corroborate the positive effects of the variant on MUC5B expression. With our models, we show that the T allele is associated with a twofold mechanism affecting *MUC5B* expression, leading to both the disruption of a DNA methylation site that directly increases *MUC5B* expression and the creation of a novel C/EBPβ transcription factor binding site that also positively regulates *MUC5B* expression.

## Material and methods

### Allele-specific match for DNA binding proteins

Motifs for DNA binding proteins were retrived through MotifDB, a bioconductor package for R, wherein we made use of the Catalog of Inferred Sequence Binding Preferences (CIS-BP) database containing information on sequence specificities for 313 human DNA binding proteins [50]. We then used matchPWM implemented in Biostrings for R to calculate PWM (position weight matrix) scores for DNA sequences with and without the *rs35705950* minor allele. P-values were assessed by scoring each of the 313 *CIS*-BP motifs on 100 thousand DNA sequences randomly selected from regulatory regions in the human genome (ChromHMM 25-state regions assigned as promoter, tss, or enhancer). We then calculate two P-values for each motif as the area under the null-curve above the observed score for the corresponding motif match against the minor and major alleles, respectively. P-values less than 0.01 were then considered as ‘candidate binding sites’ and, as an indicator of biological relevance, we considered only sites wherein the *rs35705950* minor allele alters an important nucleotide in the matching of the motifs to the DNA sequence defined here as nucleotides accounting for a mininum of 0.70 (or 70%) at a given position in the probability position matrices.

### Cell culture

The BCi_NS1.1 cell line is a human bronchial epithelial cell line kindly provided by Dr. Matthew S. Walters, Weill Cornell Medical College, New York NY, USA [51]. It was established by immortalization with retrovirus expressing human telomerase (hTERT). The bronchial epithelial cell line VA-10 was previously established by retroviral transduction of primary bronchial epithelial cells with E6 and E7 viral oncogenes [52]. Both cell lines were cultured in bronchial epithelial growth medium, BEGM (Lonza, Walkersville, MD) supplemented with 50 IU/ml penicillin and 50 μg/ml streptomycin (Gibco, Burlington, Canada).

The human lung adenocarcinoma derived alveolar epithelial cell line A549 (American Type Culture Collection, Rockville MA) was cultured in DMEM-Ham’s-F12 basal medium supplemented with 10% fetal bovine serum (FBS), 50 IU/ml penicillin and 50 μg/ml streptomycin (Gibco). All cell lines were cultured at 37°C, 5% CO_2_. All three cell lines are WT (homozygous G-allele) at *rs35705950*.

### Air-Liquid interface culture

To establish an air-liquid interface cultures (ALI), cells were seeded on the upper layer of Transwell cell culture inserts (Corning®Costar®) pore size 0.4 μm, 12 mm diameter, polyester membrane) (Sigma-Aldrich, St. Louis, USA) at density of 2×10^5^ cells per well. The cultures were maintained on chemically defined bronchial epithelial cell medium (BEGM, Cell Applications, San Diego) for 5 days, 0,5 ml in the upper chamber and 1.5 ml in the lower chamber. After 5 days, medium was changed to DMEM/F-12 (Invitrogen), supplemented with 2% UltroserG (Cergy-Saint-Christophe, France) for additional 5 days. For ALI culture, the medium was aspired from the apical side and the cell layer rinsed 1x with PBS.

### Goblet cell differentiation by IL-13 treatment

Cells were cultured for 5 days on BEGM and then for 5 days on DMEM/UG in a submerged culture. After 5 days of ALI culture, IL-13 (Peprotech, London, UK) was added to the basal side to a final concentration of 25 ng/ml and cultured for 14 days.

### Immunofluorescence staining

Cells were rinsed twice with chilled PBS. The fixation of cells was performed using 100% methanol at −20°C overnight. Subsequently, cells were submerged in 100% acetone for one minute. Staining was performed using immunofluorescence buffer, (IMF) (0.1% TX-100, 0.15M NaCL, 5nM EDTA, 20mM HEPRES, pH 7.5, 0.02% NaN3 as preservative). Cells were incubated with primary antibody overnight at +4°C, and then rinsed three times for 15 min with IMF buffer. Cells were then incubated with a secondary antibody and DAPI for two hours at room temperature, followed by four times washing with IMF buffer. Cells were mounted using ProLong Antifade (Thermo Fisher Scientific). Antibodies used for these experiments are listed in S1 Table.

### DAB staining

Paraffin-embedded tissue samples of control and IPF lung biopsies were obtained from the Department of Pathology, Landspitali University Hospital. The samples were deparaffinized, antigen retrieved by boiling in TE buffer for 20 minutes and stained with EnVisionH+ System-HRP kit (Dako) according to the manufacturer’s instructions. Primary antibodies were incubated at RT for 30 min. The immunofluorescence staining of the paraffin embedded samples (IF-P) were done as stated above with the following modifications: After antigen retrieved, samples were rinse in PBS. Primary antibody was incubated overnight. Samples were washed in PBS prior the incubation with secondary antibody and DAPI for 2 hours.

Antibodies used for these assays are listed in S1 Table. Immunofluorescence was visualized and captured using laser scanning Fluoview® FV1200 Confocal Microscope (Olympus Life Science).

### Transient transfection

Cells were grown at 70% confluence one day before transfection. FuGENE® HD Transfection Reagent (Promega) was used on BCi_NS1.1 and VA10 cells, while Lipofectamine (Thermo Fisher Scientific) was used on A549 cells. All transfection was performed following manufacturer’s instructions. The results were analysed 48h after transfection. Plasmids used for C/EBPβ overexpression was generously donated by Joan Massague: C/EBPβ LAP isoform (addgene #15738) and C/EBPβ LIP (addgene #15737) [53]. Plasmids used for this assay are listed in S2 Table.

### Production of lentiviral and cell transduction

To produce lentiviral cell lines containing pGreenFire1™ Pathway Reporter lentivector (Cat#TR010PA-N and Cat#TR000PA-1, System Bioscience) expressing the *MUC5B* promoter region and controls, we followed the general guideline provided by System Bioscience. Briefly, 70% confluent HEK-293T cells were cultured for 24 h w/o antibiotics and transfected (Lipofectamine, Thermo Fisher Scientific) with lentiviral transfection constructs and packaging plasmids (psPAX2 and pMD2.G) (Addgene plasmids #12260 and #12259, respectively). Culture medium containing the virus was harvested 24 and 48 hours post transfection and centrifuged at 1250 rpm at 4°C for 5 minutes and filtered through 0,45 μm filter. Lentiviral particle solution was added to culture medium (containing 8 μg/ml polybrene) and then added to culture flasks of 70% confluent cells (BCi_NS1.1, VA10 and A549) at a low multiplicity of infection (MOI) and incubated for 20 hours. _C_ells were then cultured further for 24 hours in fresh culture media. Infected cells were then selected with puromycin or neomycin as appropriate for 48hs. Plasmids used for this experiment are listed in S2 Table.

### Luciferase Assay

Each cell type was seeded at 70% confluence one day before transfection in a 96 well plate. To perform the luciferase assay, the Dual-Glo® Luciferase Assay System kit supplied by Promega was used, following the general guidelines provided with the kit. Luminosity was measured in a microplate reader Modulus™ II (Turner BioSystem). Luciferase measurement was normalized using Renilla co-transfection. Plasmids used for this experiment are listed in S2 Table.

### Real Time qPCR

RNA was isolated using Tri-Reagent® solution (Ambion) and cDNA preparation was carried out using RevertAid™ First strand cDNA Synthesis Kit (Fermentas) according to the manufacturer’s instructions. Real-time PCR using Power SYBR Green PCR Master mix (Applied Biosystems) was used to detect the relative quantity of each cDNA. *GAPDH* was used as the endogenous reference gene. Data were analysed using 7500 Software v2.0 (Applied Biosystems). All primers used are listed in S3 Table.

### Western Blot

Protein lysates were acquired using RIPA buffer supplemented with phosphatase and protease inhibitor cocktails (Life Technologies). For western blots, 5μg of protein was used per lane, unless otherwise stated. Samples were denatured using Laemli buffer, 10% β-mercaptoethanol at 95°C for 5 min and run on NuPage 10% Bis-Tris gels (Life Technologies) in 2-(N-morpholino) ethanesulfonic acid (MES) running buffer. Samples were then transferred to Immobilon FL PVDF membranes (Millipore). Membranes were blocked in Li-cor blocking buffer and primary antibodies were incubated overnight at 4 °C. Near-infrared fluorescence visualization was measured using Odyssey CLx scanner (Li-Cor, Cambridge, UK). Antibodies used are detailed in S1 Table.

### In Vitro Methylation Assay

DNA fragments were cut out of the pGL3-*MUC5B*pr vector using XbaI-EcoRI restriction enzymes, generating 4.1Kb of the *MUC5B* promoter region, with WT or T allele. Fragments were gel-purified using GeneJET PCR Purification Kit (ThermoFisher Scientific) following the manufacturer’s instructions and subsequently methylated with M.SssI methyltransferase (New England Biolabs) overnight at 37°C. The methylated fragments were then ligated into the pGL3 basic vector. DNA concentration were measured at 260nm before being used in transfection as described by *Vincent et al* [47]. Differential influence of methylation in the 4.1kb *MUC5B* promoter was measured by luciferase activity in three individual experiment performed in triplicates for each transfected cell line. All plasmids used are listed in S2 Table.

### Bisulfite sequencing

1×10^5^ cells were used to extract DNA from each cell line. Extraction was performed with PureLink Genomic DNA MiniKit (Invitrogen) following the manufactures instruction. Bisulfite conversion of DNA was performed with EZ DNA Methylation-Gold™ Kit (ZymoPURE™, Germany). Amplification of the region of interest was done with the EpiMark® Hot Start Taq DNA Polymerase (New England Biolab, UK) and the resulting product was sequenced by Sanger sequencing (Beckman Coulters Genomics, GENEWIZ, UK.) DNA methylation analyses of bisulfite PCR amplicons were performed using Sequence scanner V1.0. DNA methylation level was scored as percentage methylation of individual CpG units in each sample. Primers used for bisulphate sequencing are listed in S3 Table.

### 5-aza-2’-deoxycytidine DNA Methylation Inhibition

After 10 days of ALI culture, 5-aza-2’-deoxycytidine (Peprotech, London, UK) was added to basal side to a final concentration of 10μM and the cells further cultured for 3 days before being analysed. DMSO was used as a control.

### siRNA transfection

Small-interfering RNAs (siRNA) targeting human C/EBPβ were purchased from Sigma Aldrich (St Louis, MO, USA). siRNA ID: SASI_Hs02_00339146, SASI_Hs01_00236023, SASI_Hs02_00339148, SASI_Hs01_00339149, 10nM each. siRNA transfection was performed following manufactures instructions. MISSION siRNA Universal Negative Control (Sigma Aldrich) was used as a control. Cells were seeded in 96 well plates for Luciferase Assay, or in a 12 well plate to be analysed by a RT-qPCR. siRNA was transfected using Lipofectamine 2000 reagent (Invitrogen) in OPTI-MEM medium (GIBCO). Twenty-four hours later, the transfected cells were transferred to complete medium. After 48h, the cells were harvested and used for luciferase measurement and RT-qPCR.

### CRISPR

Three independent pairs of oligos targeting the *rs35705950* sequence was cloned separately into the MLM3636 guide RNA mammalian expression vector (Addgene # 43860) using the restriction enzyme BsmBI. As a control, 20 nucleotides long scrambled sgRNA was cloned into the same MLM3636 vector. gRNA vectors were corroborated by sanger sequencing. Cells were then seeded in a 12 well plate prior transfection (protocol stated above) of the spCas-9 expression vector (Addgene #44758), gRNA and homologous region and a non-homologous recombination inhibitor (Sigma-Aldrich), at 70% confluence. Selection was performed with Blasticidin. Single cell cloning assays were performed to select individual clones. Sequencing of individual clones was performed to corroborate the genotype (S3c Fig).

To generate the CRISPR pooled cells, BCi_NS1.1 and A549 cell lines selected using Blasticidin but prior single cell cloning (named pooled CRISPR cells) was used to analyze *MUC5B* expression. Sanger sequencing was done to corroborate the presence of positive edited cells in the CRISPR pool.

CRISPR control cell line was generated transfecting the spCas-9 expression vector, the scrambled gRNA and homologous region and the non-homologous recombination inhibitor. gRNA and repair template sequences are included in S3 Table.

### Statistical Analysis

Data are presented as mean and s.d. (error bars) from number of at least 3 independent experiments. Statistical differences between samples were assessed with Student two – tailed T-test. P-values below 0.05 were considered significant (***p≤0.001, **p≤0.01, *p≤.0.05).

## Results

### *Rs35705950* risk allele for IPF is associated with higher expression of MUC5B

The common polymorphism *rs35705950*, where a G is replaced by a T nucleotide, has been shown in two GWAS studies to be correlated with a predisposition to develop IPF [30, 31]. The polymorphism is located in a mucin gene cluster, three kilobases upstream of the transcription start site of *MUC5B* on chromosome 11 (S1 Fig). The molecular mechanism explaining how this polymorphism affects the aetiology of IPF has not been elucidated, although both hetero- and homozygote carriers have been shown to express higher levels of MUC5B in the lung epithelium (S1 Fig, S2 Fig) [32, 54]. To study the direct effects of this polymorphism on *MUC5B* gene regulation, a 4.1 kb of the *MUC5B cis*-regulatory region was cloned into a luciferase lentiviral reporter vector and the G of the wild type allele was replaced with the risk associated T allele using site directed mutagenesis (S3 Fig).

The resulting lentiviral vectors containing the luciferase gene under the control of 4.1kb *MUC5B* upstream region containing the promoter and either the wild type or the T variant (*rs35705950*), were used to stably transduce two human bronchial-derived basal epithelial cell lines, BCi_NS1.1 and VA10, and a human lung adenocarcinoma derived alveolar epithelial cell line A549. After stable selection, the cell lines were cultured at confluency and luciferase reporter activity measured. As shown in Fig 1a the IPF risk associated T allele consistently increased the luciferase reporter signal in all three cell lines, with the effect being most pronounced in BCi_NS1.1 cells.

**Fig 1.**
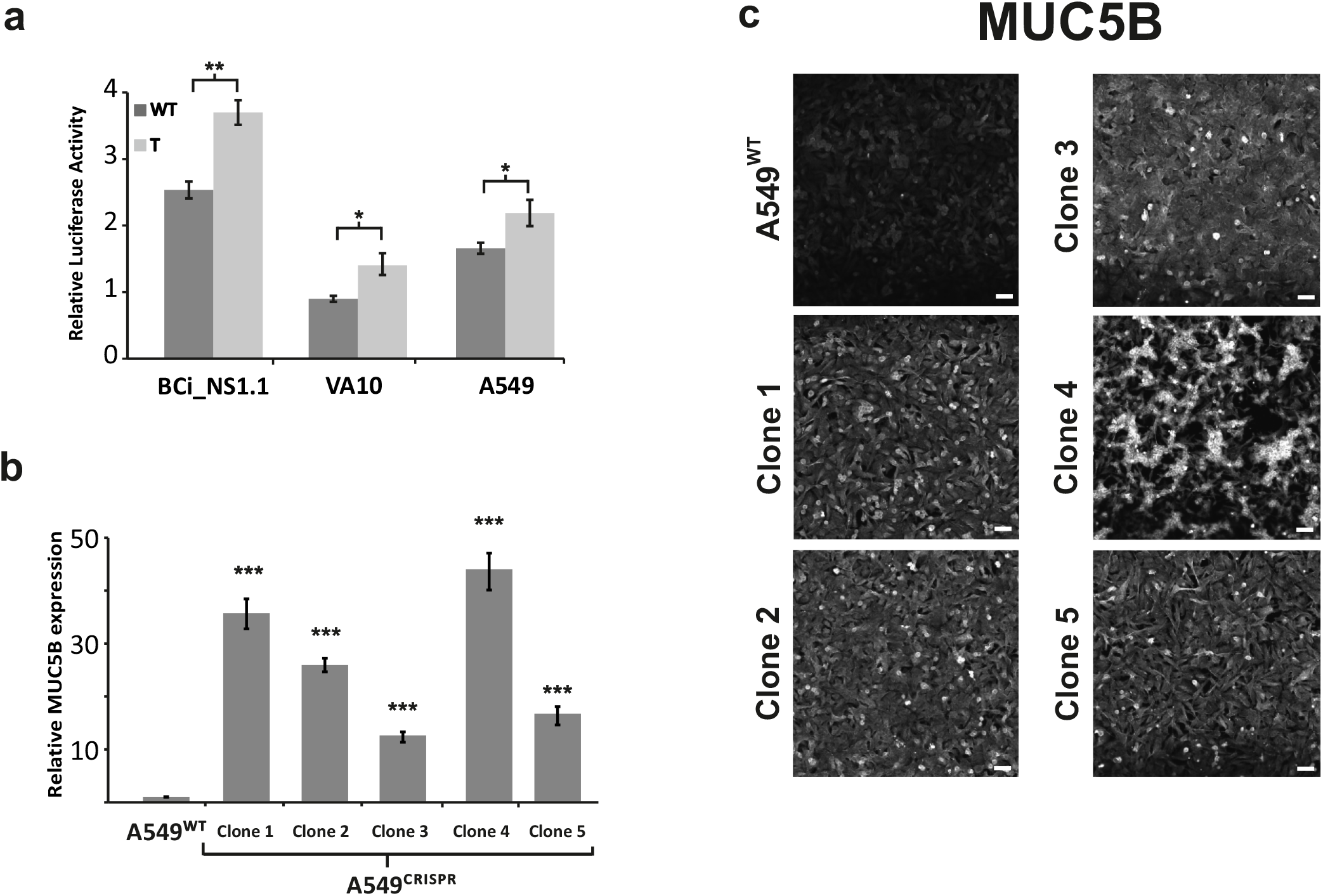
Risk allele (T) is associated with a higher expression of *MUC5B*. a) Luciferase activity on stable cell lines transduced with 4,1Kb of *MUC5B cis*-regulatory region. Luminescence was measured after 24h in a monolayer culture. b) Wild type (WT) allele was replaced on A549 cells by CRISPR editing technique to get five heterozygous cell lines [G/T]. RT-qPCR shows *MUC5B* relative expression (mRNA) in different cell lines. Culture was performed on monolayer, 48h before RNA extraction. c) IF staining on A549^CRISPR^ clones (1-5) shows higher *MUC5B* expression levels compared to A549^Cas9^ Control (WT) cell line. Scale bars = 50μm. (*p<0.05 **p<0.01 and ***p <0.001 with error bars representing SD).

To directly asses the effect of the T allele in the endogenous *rs35705950* site, we used CRISPR/Cas9 genome editing. The bronchial basal cell lines did not survive single cell cloning and thus we were only able to generate A549 edited cell lines. Five heterozygous A549^CRISPR^ clones (1-5) were generated, carrying the human [G/T] genotype (S4 Fig). The T allele had a very significant effect on the expression levels of both, MUC5B mRNA (Fig 1b) and protein (Fig 1c), showing a direct effect of the T allele on *MUC5B* expression. The above data support the previous findings [55] that the T allele confers an increased promoter/enhancer activity to the *MUC5B* upstream *cis*-regulatory sequence.

### The *rs35705950* T allele disrupts a repressive CpG DNA methylation site

It has been previously reported that the methylation of several CpG islands in the 4.1kb promoter region of *MUC5B* affects the expression of the MUC5B protein [47]. Interestingly, the presence of the T allele disrupts a CpG site that was previously characterized as methylated in the wild type. We hypothesized that the effects of the T allele could be mediated, at least in part, through epigenetic regulation. To analyse the effect of DNA methylation on *MUC5B* expression we used an air-liquid interphase (ALI) culture of BCi_NS1.1 cells. Under these conditions the cells differentiate into both, goblet cells producing mucins and ciliated cells [56]. We treated these cultures with 5’aza2’-deoxycytidine (5’AZA2’), a DNA methylation inhibitor. 5’AZA2’ treatment resulted in increased expression of *MUC5B* and *MUC5AC* indicating that DNA methylation either directly or indirectly impacts *MUC5B* expression (Fig 2a). To directly address the potential effects of DNA-methylation at the T allele, we used *M.SssI* CpG methyltransferase to *in vitro* methylate the 4.1 kb *MUC5B* promoter region. The promoter (with T allele or WT) was restriction cut from the plasmid, and the fragments was *in vitro* methylated using M.SssI CpG methyltransferase prior to ligating back into a luciferase reporter plasmid and transiently transfected into the three cell lines. We compared the effects of the mutation with (Fig 2b, bottom) or without (Fig 2b, top) direct *in vitro* methylation. In Fig 2b without *in vitro* methylation no difference is seen in luciferase activity in VA10 or A549 cells between T allele and WT while after *in vitro* methylation a significant increase is seen in the T allele carrying the luciferase plasmid compared to WT in all three cell lines. This supports the idea that differential CpG methylation of the T allele and wild type allele is at least partially involved in regulating *MUC5B* expression.

**Fig 2.**
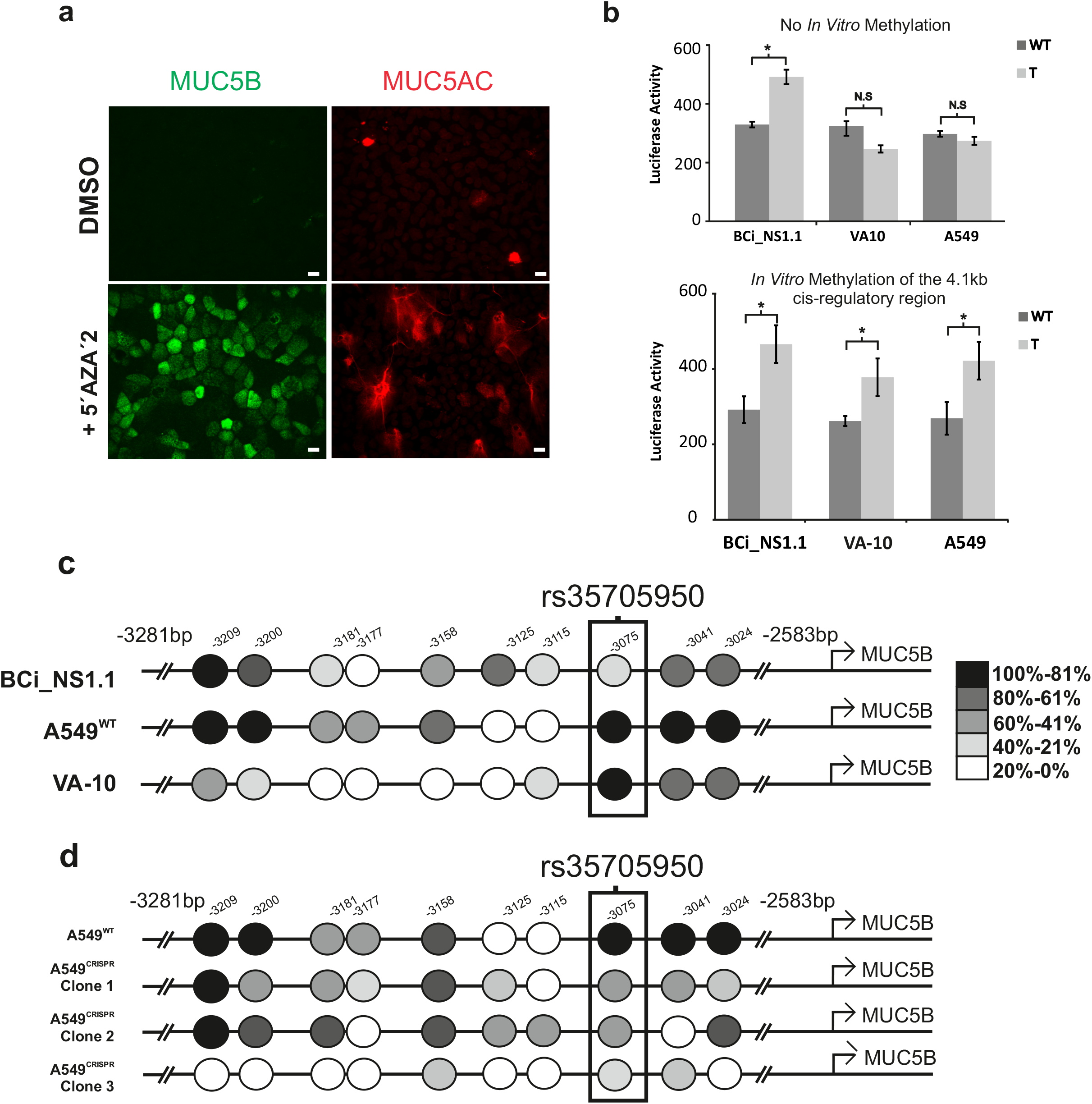
Methylation plays a role in MUC5B upregulation driven by T allele. a) IF staining shows an increased expression of MUC5B (left) and MUC5AC (right) after 48h with 5’AZA’2 treatment compare to the untreated sample under ALI conditions. Scale bar = 10μm. b) Transient transfection luciferase assay with (lower panel) or without (upper panel) *in vitro* methylation of the 4.1kb cis-regulatory region. Without in vitro methylation no difference is seen in VA10 or A549 cells between T-allele and WT while after in vitro methylation a significant increase is seen in the T-allele carrying luciferase plasmid compared to WT in all three cell lines. c) Schematic representation of a bisulfite sequencing experiment shows the differential methylation on the *rs35705950* region between cell lines (c) and how it changes with the presence of the T allele on CRISPRed cells (d). Legend indicates the range of methylation (*p<0.05 and **p <0.01 with error bars representing SD).

Targeted bisulfite sequencing of the region in our cell lines further revealed that methylation was present in all cell lines at the WT genotype although the BCi_NS1.1 cells is less methylated (21-40%), than in both VA10 and A549 cells (81-100%) under normal cell culture conditions (Fig 2d). All the cell lines are WT (G allele) homozygous carriers at the *rs35705950* site, as stated above. The CRISPR/Cas9 edited A549 cell lines harbouring the heterozygous [G/T] genotype showed less methylation (41-60%, clones 1-2, 21-40% clone 3) at the risk allele (Fig 2e) compared to A549^WT^ (81-100%), confirming that the presence of the T allele reduces methylation in the polymorphic site. This suggests that direct DNA methylation might at least partially explain the increased *MUC5B* expression in individuals carrying IPF risk associated T allele.

### C/EBPβ mediates MUC5B overexpression through *rs35705950* T allele

Even though our data support a role of differential DNA methylation to explain the effects of T allele on *MUC5B* expression, additional mechanisms might also be at play. To further study the potential effects of the T allele of *rs35705950* we used *Match*, a weight matrix-based program for predicting transcription factor binding sites in DNA sequences using the DNA sequence flanking *rs35705950* polymorphism. The program uses a library of positional weight matrices from TRANSFAC® Public 6.0. Fig 3a shows the predicted transcription factor binding motif. Binding motifs for PAX2 and PAX4 overlap the polymorphism in the WT sequence while the T allele leads to the loss of the PAX4 motif. The T allele furthermore leads to a gain of a novel *CCAAT*/enhancer-binding proteins (C/EBPs) motif (Fig 3a, S6a,S6b Fig, S3 Table). Due to the low expression levels of PAX2 and PAX4 in lung tissue (S2b Fig) they are not likely to be causative of the T allele increased promoter activity. On the other hand, in the C/EBP family, C/EBPβ has a documented role in inflammation [57, 58] and C/EBPβ is highly expressed in lung tissue (S2a Fig), making it an ideal candidate to further studies. As shown in S2c Fig, the eQTL effects of the *rs35705950* polymorphism is primarily seen in the lung, suggesting lung specific effects. To further address whether the T allele might lead to a novel C/EBPβ binding site we carried out a direct comparison of a predicted C/EBPβ binding using motifbreakR [59]. S6a Fig shows that there is a predicted consensus binding site for C/EBPβ on the negative strand in the minor allele. On the negative strand the T-allele is now A and the WT allele a C. The PWM (position weight matrix) score for the A allele is 89% (P = 0.0041) compared to a PWM score of 71% (P = 0.103) for the C allele (the major allele), novel binding of C/EBPβ to the negative strand of the IPF risk allele.

**Fig 3.**
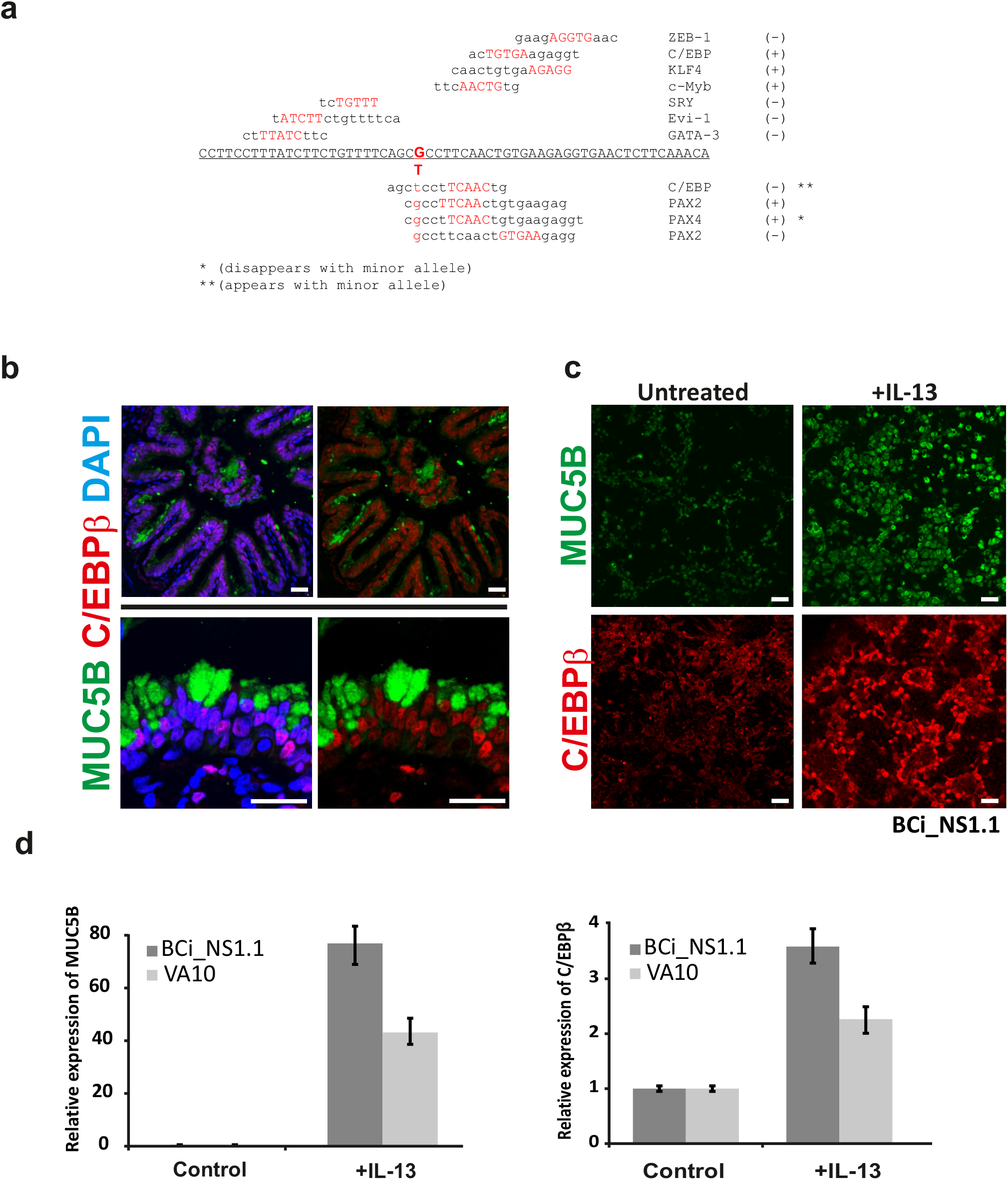
C/EBP is predicted to bind rs35705950 MUC5B cis-regulatory region only in presence of the T allele and C/EBPβ is co-expressed with MUC5B in IPF. a) Results of a weight matrix-based program (Match) for predicting transcription factor binding sites in *MUC5B* promoter sequence using the DNA sequence flanking *rs35705950* using TRANSFAC©. ** indicates a novel binding motif in the T-variant allele. * indicates the loss of a transcription factor binding motif, while (+) and (−) indicates the strand where the bindings occurs. b) IF-P in an IPF sample shows the co-expression of MUC5B (green) and C/EBPβ (red). c) IL-13 (20ng/mL) increases MUC5B (green) and C/EBPβ (red) expression at protein (c) and at RNA level (d), analysed by RT-qPCR, compared to the untreated sample. Scale bars = 50μm.

C/EBPβ has three isoforms generated by alternative splicing (S1 Fig). Two of these isoforms (LAP and LAP*) mediate transcriptional activation through the transactivation domain, while the third isoform (LIP) has an inhibitory role due to the lack of the transactivation domain [60].

Immunofluorescence staining on four IPF lung samples show that C/EBPβ and MUC5B are co-expressed in airway epithelial cells (Fig 3b, S3 Fig). Furthermore, in healthy lung samples C/EBPβ is highly expressed in alveolar cells resembling alveolar macrophages and they also express high levels of MUC5B (S3 Fig).

In addition to the aforementioned co-expression, C/EBPβ might act through a common signalling pathway potentially driving *MUC5B* overexpression through the T allele. IL-13 has been previously used to induce goblet cell hyperplasia in asthma models and has also been used to induce *MUC5AC* expression in cell culture models [56]. We cultured both VA10 and BCi_NS1.1 cell lines, under ALI culture conditions with or without IL-13. In both cell lines, IL-13 induced C/EBPβ as well as *MUC5B* expression at both the protein (Fig 3c) and mRNA level (Fig 3d), suggesting a co-regulatory pathway.

To analyse specifically the differential effects of C/EBPβ on *MUC5B cis*-regulatory domain on the wild type and T alleles, we overexpressed the C/EBPβ isoforms in the BCi_NS1.1 (Fig 4a), A549 (Fig 4b) and VA10 (S6c Fig) cell lines harbouring the stably integrated luciferase reporter associated with the 4.1kb *MUC5B* promoter region. The C/EBPβ LAP* isoform increased *MUC5B* promoter regulated luciferase activity 2.5-fold when the T allele was present, compared to the wild type in both BCi_NS1.1 and A549. This relationship is dose-dependent, as shown in S6d Fig. Overexpression of the C/EBPβ LIP isoform reduced *MUC5B* reporter activity when the T allele was present in both cell lines (Fig 4 a, b). When both isoforms were co-expressed in equimolar concentrations an increased luciferase signal was seen in the BCi_NS1.1 cell line.

**Fig 4.**
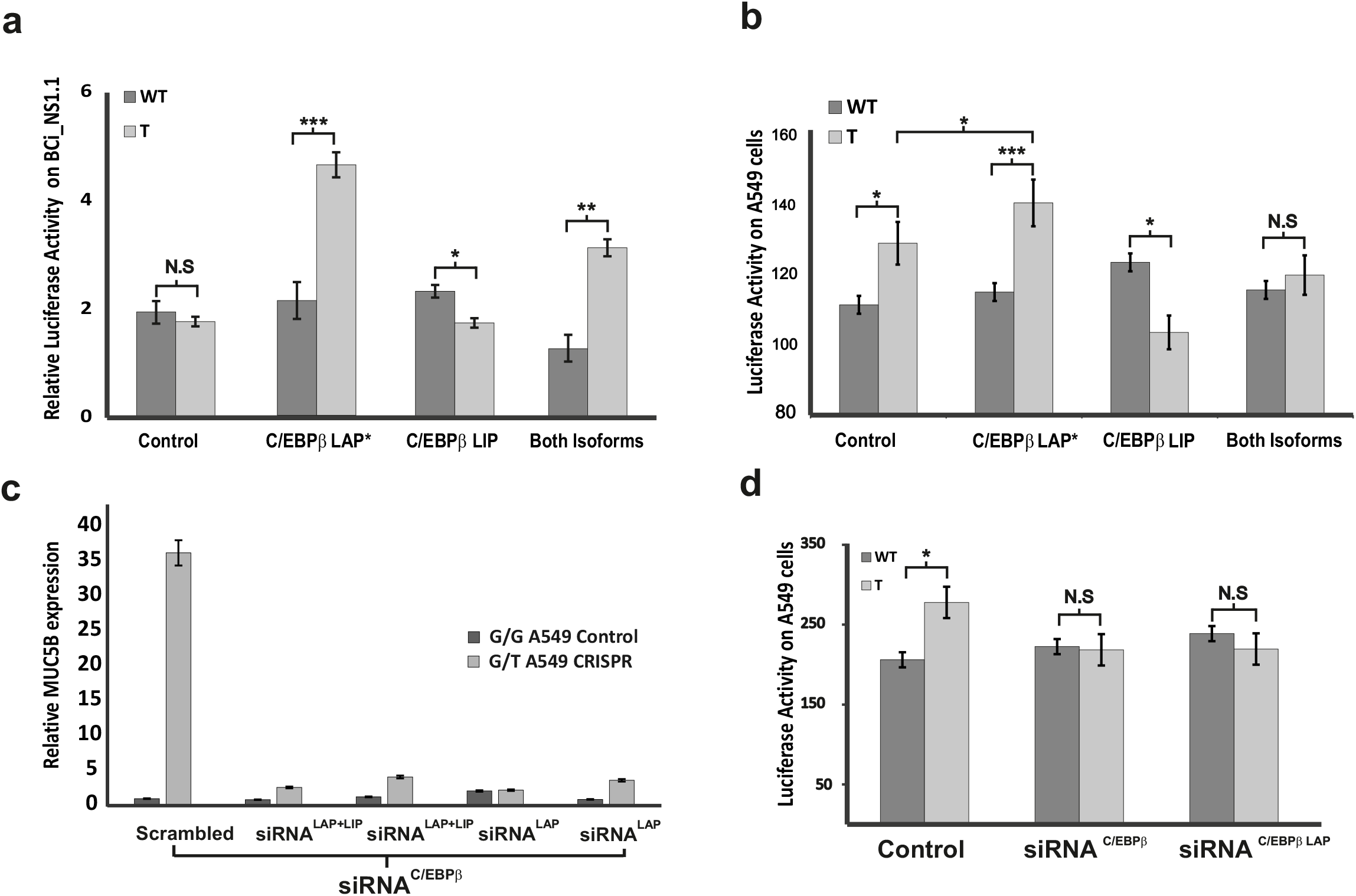
C/EBPβ mediates the MUC5B upregulation by the rs35705950 T allele. a) The three C/EBPβ isoforms were transfected individually into BCi_NS1.1 (a) and A549 (b) luciferase stable cell lines carrying the 4.1kb *MUC5B* promoter for 24h. The activatory isoform (LAP) increases the *MUC5B* reporter activity only in presence of the T allele, while the inhibitory isoform (LIP) inhibits the differential activity between WT and T allele. c) siRNA silencing C/EBPβ isoforms, LAP and LIP together or C/EBPβ LAP alone, was performed on A549^CRISPR^ Clone 1 compared to A549^Cas9^ Control cell line. Inhibition of C/EBPβ restores differential *MUC5B* expression driven by the T allele to the WT associated *MUC5B* expression levels while no effect is seen in A549^Cas9^ Control cell line. d) siRNA was used to knowck down C/EBPβ (LAP and LIP or only LAP isoform) in A549 luciferase stable cell line carrying the 4.1kb *MUC5B* promoter. Inhibition of C/EBPβ restores *MUC5B* expression driven by the T allele to the WT associated *MUC5B* expression levels (*p<0.05 and **p <0.01 with error bars representing SD).

To further corroborate the role of C/EBPβ in regulating *MUC5B* expression through the risk allele, we knocked down C/EBPβ expression using siRNAs targeting either the LAP/LAP* or all isoforms. Stable cell lines carrying the luciferase reporter vector and A549^CRISPR^ clones were used to analyse the effect of C/EBPβ knock-down on *MUC5B* expression by RT-qPCR (Fig 4c, S6e Fig) and luciferase reporter assay (Fig 4d, S6f Fig). Our results show that siRNAs targeting C/EBPβ lowers *MUC5B* expression in T allele genotype to a similar level as wild type.

The correlation between C/EBPβ expression and the strong T allele specific positive expression suggests that this transcription factor is a candidate key regulator of *MUC5B* expression in carriers of the T allele in IPF.

## Discussion

The results presented in this study suggest that the IPF associated polymorphism, *rs35705950* (T allele), has a direct and major effect on *MUC5B* expression. We have shown that it is mediated through a combination of effects on both epigenetic and transcriptional regulatory mechanisms. Specifically, we have shown that the T allele directly reduces DNA methylation and that this increases the *MUC5B* expression in our cell culture models. Furthermore, we have shown that the T allele creates a novel C/EBPβ binding site that strongly positively regulates *MUC5B* expression.

CpG DNA methylation has previously been identified as an important regulatory mechanism for mucins. The human genes *MUC2*, *MUC5AC*, *MUC5B* and *MUC6* are clustered on chromosome *11p15* and their promoters show a GC-rich structure [47]. Hypermethylation of *MUC5B* promoter is the major mechanism responsible for its silencing, combined with histone acetylation while the expression of *MUC5AC* was rarely influenced by epigenetic marks. In this study, we confirm a general role of methylation in *MUC5B* repression. Furthermore, we show that the *rs35705950* T allele disrupts a CpG methylation site and this disruption relieves a DNA methylation-dependent negative regulation site. Furthermore, the *in vitro* methylation assay corroborates the differential methylation pattern between alleles.

Data from the ENCODE project suggest that the area surrounding the *MUC5B* polymorphism is located within a *cis*-regulatory element, based on both, the DNAse hypersensitivity cluster and the high number of transcription factors binding in the area. Our data, based on *in silico* binding analysis, predicts a C/EBPβ binding site overlapping *rs35705950* when the IPF associated T allele is present. Here, we directly show that *rs35705950* polymorphism leads to a specific response to C/EBPβ positively regulating MUC5B expression. C/EBPβ would thus be implicated as an important regulator of MUC5B expression in carriers of the *rs35705950* T allele.

Mucin regulating signalling pathways are complex and poorly understood. It has been shown that activation of MEK1/2, PI3K, SPhk1, and MAPK14 (p38α-MAPK) are implicated in IL-13 induced mucus production [61, 62]. Furthermore, through PMA signalling PKC, EGF/TGF-a, Ras/Raf, Mek, ERK and Sp-1 signalling pathways have also been associated with *MUC5AC* and *MUC5B* expression [63]. Interestingly, C/EBPβ is also overexpressed after PMA stimulus [64] and, in our results, we show a common signalling pathway between MUC5B and C/EBPβ involving IL-13, suggesting common signalling pathways. C/EBPβ has been previously associated with inflammation and immune responses [58, 65–67]. Similarly, MUC5B has been implicated in innate immune response in the lung [68], although C/EBPβ has not been implicated so far. C/EBPβ is also expressed in macrophages [69, 70]. Recently, Satoh *et al*.[71] identified a monocyte derived cell, termed segregated-nucleus-containing atypical monocyte (SatM). These SatM cells were shown to be crucial for bleomycin induced fibrosis in a mouse model and C/EBPβ was shown to be a key transcriptional regulator in their differentiation. Interestingly, the SatM-termed monocytes, induce a pro-fibrotic signalling pathway linking them directly to the fibrosis. Furthermore, macrophages have been shown to be related to goblet cell hyperplasia and in that context to induce *MUC5B* but not *MUC5AC* in human bronchial epithelial cells [72]. Whether macrophages play any role in dysregulation of *MUC5B* in IPF remains to be tested.

Other potential transcriptional binding sites are present in the region of the *rs35705950* polymorphic site. Highly preserved HOX9 and a FOXA2 binding domains have been described [55, 73]. The binding of FOXA2 transcription factor was shown to be 32bp upstream of *rs35705950* polymorphism. The proximity to the polymorphism may suggest a co-regulatory site that could through, e.g. interaction with the novel C/EBPβ binding site affect *MUC5B* expression. Whether the overexpression is caused by a combination of FOXA2 and C/EBPβ needs to be clarified. The eQTL data showing that the positive expression effects of the *rs35705950* polymorphic site is primarily seen in the lung (S2d Fig) could be related to a lung specific pattern of expression of transcription factors, e.g. C/EBPβ with or without other coregulators such as FOXA2. Other tissues that express high levels of MUC5B, such as salivary glands, stomach, small and large intestine are not shown to have a positive eQTL with the polymorphic site.

In our results, we have also shown that C/EBPβ is co-expressed with MUC5B. Its presence is predominantly in basal cells in the pseudostratified epithelium under normal conditions, while in fibrotic lungs, it is also expressed in ciliated and goblet cells. This could be related with the role of C/EBPβ previously associated with fibrosis development [74, 75].

To summarize, we have shown that T allele associated with the *rs35705950* polymorphism strongly induces *MUC5B* overexpression. Interestingly, this appears to be through two independent mechanisms. Firstly, it leads to disruption of a CpG methylation site that naturally represses *MUC5B*, resulting in an overexpression. Secondly, the same T allele creates a potential novel C/EBPβ binding site that positively regulates *MUC5B* expression. These results identify the C/EBPβ transcription factor as a candidate target to study in fibrosis associated with IPF. Further studies are needed to decipher how C/EBPβ and MUC5B can result in fibrosis.

## Acknowledgements

We would like to thank members of the Stem Cell Research Unit, especially Sævar Ingþórsson and Bryndis Valdimarsdottir for various help with cell culture and staining. We would like to thank Helgi Isaksson, MD for providing IPF samples.

## Declarations

### Ethics approval and consent to participate

Not applicable.

### Consent for publication

Not applicable.

### Availability of data and material

The datasets used and/or analysed during the current study are available from the corresponding author on request.

### Authors’ contributions

Conception and design: AUA, MKM;

Analysis and interpretation: AUA, AJA, OAS, GG, TG, MKM;

Drafting the manuscript for important intellectual content: AUA, MKM

## References

1. Hutchinson JP, McKeever TM, Fogarty AW, Navaratnam V, Hubbard RB. Increasing global mortality from idiopathic pulmonary fibrosis in the twenty-first century. Ann Am Thorac Soc. 2014;11(8):1176–85. Epub 2014/08/29. doi: 10.1513/AnnalsATS.201404-145OC. PubMed PMID: 25165873.

2. Coultas DB, Zumwalt RE, Black WC, Sobonya RE. The epidemiology of interstitial lung diseases. Am J Respir Crit Care Med. 1994;150(4):967–72. Epub 1994/10/01. doi: 10.1164/ajrccm.150.4.7921471. PubMed PMID: 7921471.

3. Hirakawa H, Pierce RA, Bingol-Karakoc G, Karaaslan C, Weng M, Shi GP, et al. Cathepsin S deficiency confers protection from neonatal hyperoxia-induced lung injury. Am J Respir Crit Care Med. 2007;176(8):778–85. Epub 2007/08/04. doi: 10.1164/rccm.200704-519OC. PubMed PMID: 17673697; PubMed Central PMCID: PMCPMC2020827.

4. Hunninghake GM, Hatabu H, Okajima Y, Gao W, Dupuis J, Latourelle JC, et al. MUC5B promoter polymorphism and interstitial lung abnormalities. N Engl J Med. 2013;368(23):2192–200. Epub 2013/05/23. doi: 10.1056/NEJMoa1216076. PubMed PMID: 23692170; PubMed Central PMCID: PMCPMC3747636.

5. Olson AL, Swigris JJ, Lezotte DC, Norris JM, Wilson CG, Brown KK. Mortality from pulmonary fibrosis increased in the United States from 1992 to 2003. Am J Respir Crit Care Med. 2007;176(3):277–84. Epub 2007/05/05. doi: 10.1164/rccm.200701-044OC. PubMed PMID: 17478620.

6. Raghu G, Collard HR, Egan JJ, Martinez FJ, Behr J, Brown KK, et al. An official ATS/ERS/JRS/ALAT statement: idiopathic pulmonary fibrosis: evidence-based guidelines for diagnosis and management. Am J Respir Crit Care Med. 2011;183(6):788–824. Epub 2011/04/08. doi: 10.1164/rccm.2009-040GL. PubMed PMID: 21471066; PubMed Central PMCID: PMCPMC5450933.

7. Margaritopoulos GA, Vasarmidi E, Antoniou KM. Pirfenidone in the treatment of idiopathic pulmonary fibrosis: an evidence-based review of its place in therapy. Core Evid. 2016;11:11–22. Epub 2016/07/23. doi: 10.2147/CE.S76549. PubMed PMID: 27445644; PubMed Central PMCID: PMCPMC4936814.

8. Bonella F, Stowasser S, Wollin L. Idiopathic pulmonary fibrosis: current treatment options and critical appraisal of nintedanib. Drug Des Devel Ther. 2015;9:6407–19. Epub 2015/12/31. doi: 10.2147/DDDT.S76648. PubMed PMID: 26715838; PubMed Central PMCID: PMCPMC4686227.

9. Hubbard R, Cooper M, Antoniak M, Venn A, Khan S, Johnston I, et al. Risk of cryptogenic fibrosing alveolitis in metal workers. Lancet. 2000;355(9202):466–7. Epub 2000/06/07. doi: 10.1016/S0140-6736(00)82017-6. PubMed PMID: 10841131.

10. Hubbard R, Lewis S, Richards K, Johnston I, Britton J. Occupational exposure to metal or wood dust and aetiology of cryptogenic fibrosing alveolitis. Lancet. 1996;347(8997):284–9. Epub 1996/02/03. PubMed PMID: 8569361.

11. Iwai K, Mori T, Yamada N, Yamaguchi M, Hosoda Y. Idiopathic pulmonary fibrosis. Epidemiologic approaches to occupational exposure. Am J Respir Crit Care Med. 1994;150(3):670–5. Epub 1994/09/01. doi: 10.1164/ajrccm.150.3.8087336. PubMed PMID: 8087336.

12. Baumgartner KB, Samet JM, Coultas DB, Stidley CA, Hunt WC, Colby TV, et al. Occupational and environmental risk factors for idiopathic pulmonary fibrosis: a multicenter case-control study. Collaborating Centers. Am J Epidemiol. 2000;152(4):307–15. Epub 2000/09/01. PubMed PMID: 10968375.

13. Lawson WE, Crossno PF, Polosukhin VV, Roldan J, Cheng DS, Lane KB, et al. Endoplasmic reticulum stress in alveolar epithelial cells is prominent in IPF: association with altered surfactant protein processing and herpesvirus infection. Am J Physiol Lung Cell Mol Physiol. 2008;294(6):L1119–26. Epub 2008/04/09. doi: 10.1152/ajplung.00382.2007. PubMed PMID: 18390830.

14. Stewart JP, Egan JJ, Ross AJ, Kelly BG, Lok SS, Hasleton PS, et al. The detection of Epstein-Barr virus DNA in lung tissue from patients with idiopathic pulmonary fibrosis. Am J Respir Crit Care Med. 1999;159(4 Pt 1):1336–41. Epub 1999/04/08. doi: 10.1164/ajrccm.159.4.9807077. PubMed PMID: 10194186.

15. Tang YW, Johnson JE, Browning PJ, Cruz-Gervis RA, Davis A, Graham BS, et al. Herpesvirus DNA is consistently detected in lungs of patients with idiopathic pulmonary fibrosis. J Clin Microbiol. 2003;41(6):2633–40. Epub 2003/06/07. PubMed PMID: 12791891; PubMed Central PMCID: PMCPMC156536.

16. Erwteman TM, Braat MC, van Aken WG. Interstitial pulmonary fibrosis: a new side effect of practolol. Br Med J. 1977;2(6082):297–8. Epub 1977/07/30. PubMed PMID: 871866; PubMed Central PMCID: PMCPMC1630827.

17. Hubbard R, Venn A, Smith C, Cooper M, Johnston I, Britton J. Exposure to commonly prescribed drugs and the etiology of cryptogenic fibrosing alveolitis: a case-control study. Am J Respir Crit Care Med. 1998;157(3 Pt 1):743–7. Epub 1998/03/28. doi: 10.1164/ajrccm.157.3.9701093. PubMed PMID: 9517585.

18. Musk AW, Pollard JA. Pindolol and pulmonary fibrosis. Br Med J. 1979;2(6190):581–2. Epub 1979/09/08. PubMed PMID: 497711; PubMed Central PMCID: PMCPMC1596501.

19. Baumgartner KB, Samet JM, Stidley CA, Colby TV, Waldron JA. Cigarette smoking: a risk factor for idiopathic pulmonary fibrosis. Am J Respir Crit Care Med. 1997;155(1):242–8. Epub 1997/01/01. doi: 10.1164/ajrccm.155.1.9001319. PubMed PMID: 9001319.

20. Spira A, Beane J, Shah V, Liu G, Schembri F, Yang X, et al. Effects of cigarette smoke on the human airway epithelial cell transcriptome. Proc Natl Acad Sci U S A. 2004;101(27):10143–8. Epub 2004/06/24. doi: 10.1073/pnas.0401422101. PubMed PMID: 15210990; PubMed Central PMCID: PMCPMC454179.

21. Steele MP, Speer MC, Loyd JE, Brown KK, Herron A, Slifer SH, et al. Clinical and pathologic features of familial interstitial pneumonia. Am J Respir Crit Care Med. 2005;172(9):1146–52. Epub 2005/08/20. doi: 10.1164/rccm.200408-1104OC. PubMed PMID: 16109978; PubMed Central PMCID: PMCPMC2718398.

22. Tsakiri KD, Cronkhite JT, Kuan PJ, Xing C, Raghu G, Weissler JC, et al. Adult-onset pulmonary fibrosis caused by mutations in telomerase. Proc Natl Acad Sci U S A. 2007;104(18):7552–7. Epub 2007/04/27. doi: 10.1073/pnas.0701009104. PubMed PMID: 17460043; PubMed Central PMCID: PMCPMC1855917.

23. Armanios MY, Chen JJ, Cogan JD, Alder JK, Ingersoll RG, Markin C, et al. Telomerase mutations in families with idiopathic pulmonary fibrosis. N Engl J Med. 2007;356(13):1317–26. Epub 2007/03/30. doi: 10.1056/NEJMoa066157. PubMed PMID: 17392301.

24. Cogan JD, Kropski JA, Zhao M, Mitchell DB, Rives L, Markin C, et al. Rare variants in RTEL1 are associated with familial interstitial pneumonia. Am J Respir Crit Care Med. 2015;191(6):646–55. Epub 2015/01/22. doi: 10.1164/rccm.201408-1510OC. PubMed PMID: 25607374; PubMed Central PMCID: PMCPMC4384777.

25. Stuart BD, Choi J, Zaidi S, Xing C, Holohan B, Chen R, et al. Exome sequencing links mutations in PARN and RTEL1 with familial pulmonary fibrosis and telomere shortening. Nat Genet. 2015;47(5):512–7. Epub 2015/04/08. doi: 10.1038/ng.3278. PubMed PMID: 25848748; PubMed Central PMCID: PMCPMC4414891.

26. Thomas AQ, Lane K, Phillips J, 3rd, Prince M, Markin C, Speer M, et al. Heterozygosity for a surfactant protein C gene mutation associated with usual interstitial pneumonitis and cellular nonspecific interstitial pneumonitis in one kindred. Am J Respir Crit Care Med. 2002;165(9):1322–8. Epub 2002/05/07. doi: 10.1164/rccm.200112-123OC. PubMed PMID: 11991887.

27. van Moorsel CH, van Oosterhout MF, Barlo NP, de Jong PA, van der Vis JJ, Ruven HJ, et al. Surfactant protein C mutations are the basis of a significant portion of adult familial pulmonary fibrosis in a dutch cohort. Am J Respir Crit Care Med. 2010;182(11):1419–25. Epub 2010/07/27. doi: 10.1164/rccm.200906-0953OC. PubMed PMID: 20656946.

28. Wang Y, Kuan PJ, Xing C, Cronkhite JT, Torres F, Rosenblatt RL, et al. Genetic defects in surfactant protein A2 are associated with pulmonary fibrosis and lung cancer. Am J Hum Genet. 2009;84(1):52–9. Epub 2008/12/23. doi: 10.1016/j.ajhg.2008.11.010. PubMed PMID: 19100526; PubMed Central PMCID: PMCPMC2668050.

29. Kropski JA, Lawson WE, Young LR, Blackwell TS. Genetic studies provide clues on the pathogenesis of idiopathic pulmonary fibrosis. Dis Model Mech. 2013;6(1):9–17. Epub 2012/12/27. doi: 10.1242/dmm.010736. PubMed PMID: 23268535; PubMed Central PMCID: PMCPMC3529334.

30. Noth I, Zhang YZ, Ma SF, Flores C, Barber M, Huang Y, et al. Genetic variants associated with idiopathic pulmonary fibrosis susceptibility and mortality: a genome-wide association study. Lancet Resp Med. 2013;1(4):309–17. doi: 10.1016/S2213-2600(13)70045-6. PubMed PMID: WOS:000342689000018.

31. Fingerlin TE, Murphy E, Zhang W, Peljto AL, Brown KK, Steele MP, et al. Genome-wide association study identifies multiple susceptibility loci for pulmonary fibrosis. Nat Genet. 2013;45(6):613–20. Epub 2013/04/16. doi: 10.1038/ng.2609. PubMed PMID: 23583980; PubMed Central PMCID: PMCPMC3677861.

32. Seibold MA, Wise AL, Speer MC, Steele MP, Brown KK, Loyd JE, et al. A common MUC5B promoter polymorphism and pulmonary fibrosis. N Engl J Med. 2011;364(16):1503–12. Epub 2011/04/22. doi: 10.1056/NEJMoa1013660. PubMed PMID: 21506741; PubMed Central PMCID: PMCPMC3379886.

33. Evans CM, Fingerlin TE, Schwarz MI, Lynch D, Kurche J, Warg L, et al. Idiopathic Pulmonary Fibrosis: A Genetic Disease That Involves Mucociliary Dysfunction of the Peripheral Airways. Physiol Rev. 2016;96(4):1567–91. Epub 2016/09/16. doi: 10.1152/physrev.00004.2016. PubMed PMID: 27630174; PubMed Central PMCID: PMCPMC5243224.

34. Praetorius C, Grill C, Stacey SN, Metcalf AM, Gorkin DU, Robinson KC, et al. A polymorphism in IRF4 affects human pigmentation through a tyrosinase-dependent MITF/TFAP2A pathway. Cell 2013;155(5):1022–33. Epub 2013/11/26. doi: 10.1016/j.cell.2013.10.022. PubMed PMID: 24267888; PubMed Central PMCID: PMCPMC3873608.

35. Gupta RM, Hadaya J, Trehan A, Zekavat SM, Roselli C, Klarin D, et al. A Genetic Variant Associated with Five Vascular Diseases Is a Distal Regulator of Endothelin-1 Gene Expression. Cell 2017;170(3):522–33 e15. Epub 2017/07/29. doi: 10.1016/j.cell.2017.06.049. PubMed PMID: 28753427.

36. Tak YG, Farnham PJ. Making sense of GWAS: using epigenomics and genome engineering to understand the functional relevance of SNPs in non-coding regions of the human genome. Epigenetics Chromatin. 2015;8:57. Epub 2016/01/01. doi: 10.1186/s13072-015-0050-4. PubMed PMID: 26719772; PubMed Central PMCID: PMCPMC4696349.

37. Borie R, Crestani B, Dieude P, Nunes H, Allanore Y, Kannengiesser C, et al. The MUC5B variant is associated with idiopathic pulmonary fibrosis but not with systemic sclerosis interstitial lung disease in the European Caucasian population. PLoS One. 2013;8(8):e70621. Epub 2013/08/14. doi: 10.1371/journal.pone.0070621. PubMed PMID: 23940607; PubMed Central PMCID: PMCPMC3734256.

38. Horimasu Y, Ohshimo S, Bonella F, Tanaka S, Ishikawa N, Hattori N, et al. MUC5B promoter polymorphism in Japanese patients with idiopathic pulmonary fibrosis. Respirology. 2015;20(3):439–44. Epub 2015/01/13. doi: 10.1111/resp.12466. PubMed PMID: 25581455.

39. Stock CJ, Sato H, Fonseca C, Banya WA, Molyneaux PL, Adamali H, et al. Mucin 5B promoter polymorphism is associated with idiopathic pulmonary fibrosis but not with development of lung fibrosis in systemic sclerosis or sarcoidosis. Thorax. 2013;68(5):436–41. Epub 2013/01/17. doi: 10.1136/thoraxjnl-2012-201786. PubMed PMID: 23321605.

40. Wei R, Li C, Zhang M, Jones-Hall YL, Myers JL, Noth I, et al. Association between MUC5B and TERT polymorphisms and different interstitial lung disease phenotypes. Transl Res. 2014;163(5):494–502. Epub 2014/01/18. doi: 10.1016/j.trsl.2013.12.006. PubMed PMID: 24434656; PubMed Central PMCID: PMCPMC4074379.

41. Zhang Y, Noth I, Garcia JG, Kaminski N. A variant in the promoter of MUC5B and idiopathic pulmonary fibrosis. N Engl J Med. 2011;364(16):1576–7. Epub 2011/04/22. doi: 10.1056/NEJMc1013504. PubMed PMID: 21506748; PubMed Central PMCID: PMCPMC4327944.

42. Peljto AL, Selman M, Kim DS, Murphy E, Tucker L, Pardo A, et al. The MUC5B promoter polymorphism is associated with idiopathic pulmonary fibrosis in a Mexican cohort but is rare among Asian ancestries. Chest. 2015;147(2):460–4. Epub 2014/10/03. doi: 10.1378/chest.14-0867. PubMed PMID: 25275363; PubMed Central PMCID: PMCPMC4314820.

43. Wang C, Zhuang Y, Guo W, Cao L, Zhang H, Xu L, et al. Mucin 5B promoter polymorphism is associated with susceptibility to interstitial lung diseases in Chinese males. PLoS One. 2014;9(8):e104919. Epub 2014/08/15. doi: 10.1371/journal.pone.0104919. PubMed PMID: 25121989; PubMed Central PMCID: PMCPMC4133265.

44. Genomes Project C, Auton A, Brooks LD, Durbin RM, Garrison EP, Kang HM, et al. A global reference for human genetic variation. Nature. 2015;526(7571):68–74. Epub 2015/10/04. doi: 10.1038/nature15393. PubMed PMID: 26432245; PubMed Central PMCID: PMCPMC4750478.

45. Roy MG, Rahmani M, Hernandez JR, Alexander SN, Ehre C, Ho SB, et al. Mucin production during prenatal and postnatal murine lung development. Am J Respir Cell Mol Biol. 2011;44(6):755–60. Epub 2011/06/10. doi: 10.1165/rcmb.2010-0020OC 10.1165/rcmb.2010-0020RC. PubMed PMID: 21653907; PubMed Central PMCID: PMCPMC3135838.

46. Young HW, Williams OW, Chandra D, Bellinghausen LK, Perez G, Suarez A, et al. Central role of Muc5ac expression in mucous metaplasia and its regulation by conserved 5’ elements. Am J Respir Cell Mol Biol. 2007;37(3):273–90. Epub 2007/04/28. doi: 10.1165/rcmb.2005-0460OC. PubMed PMID: 17463395; PubMed Central PMCID: PMCPMC1994232.

47. Vincent A, Perrais M, Desseyn JL, Aubert JP, Pigny P, Van Seuningen I. Epigenetic regulation (DNA methylation, histone modifications) of the 11p15 mucin genes (MUC2, MUC5AC, MUC5B, MUC6) in epithelial cancer cells. Oncogene. 2007;26(45):6566–76. Epub 2007/05/02. doi: 10.1038/sj.onc.1210479. PubMed PMID: 17471237.

48. Heazlewood CK, Cook MC, Eri R, Price GR, Tauro SB, Taupin D, et al. Aberrant mucin assembly in mice causes endoplasmic reticulum stress and spontaneous inflammation resembling ulcerative colitis. PLoS Med. 2008;5(3):e54. Epub 2008/03/06. doi: 10.1371/journal.pmed.0050054. PubMed PMID: 18318598; PubMed Central PMCID: PMCPMC2270292.

49. Romero F, Summer R. Protein Folding and the Challenges of Maintaining Endoplasmic Reticulum Proteostasis in Idiopathic Pulmonary Fibrosis. Ann Am Thorac Soc. 2017;14(Supplement_5):S410–S3. Epub 2017/11/22. doi: 10.1513/AnnalsATS.201703-207AW. PubMed PMID: 29161089; PubMed Central PMCID: PMCPMC5711273.

50. Weirauch MT, Yang A, Albu M, Cote AG, Montenegro-Montero A, Drewe P, et al. Determination and inference of eukaryotic transcription factor sequence specificity. Cell. 2014;158(6):1431–43. Epub 2014/09/13. doi: 10.1016/j.cell.2014.08.009. PubMed PMID: 25215497; PubMed Central PMCID: PMCPMC4163041.

51. Walters MS, Gomi K, Ashbridge B, Moore MA, Arbelaez V, Heldrich J, et al. Generation of a human airway epithelium derived basal cell line with multipotent differentiation capacity. Respir Res. 2013;14:135. Epub 2013/12/05. doi: 10.1186/1465-9921-14-135. PubMed PMID: 24298994; PubMed Central PMCID: PMCPMC3907041.

52. Halldorsson S, Asgrimsson V, Axelsson I, Gudmundsson GH, Steinarsdottir M, Baldursson O, et al. Differentiation potential of a basal epithelial cell line established from human bronchial explant. In Vitro Cell Dev Biol Anim. 2007;43(8-9):283–9. Epub 2007/09/19. doi: 10.1007/s11626-007-9050-4. PubMed PMID: 17876679.

53. Gomis RR, Alarcon C, Nadal C, Van Poznak C, Massague J. C/EBPbeta at the core of the TGFbeta cytostatic response and its evasion in metastatic breast cancer cells. Cancer Cell. 2006;10(3):203–14. Epub 2006/09/09. doi: 10.1016/j.ccr.2006.07.019. PubMed PMID: 16959612.

54. Seibold MA, Smith RW, Urbanek C, Groshong SD, Cosgrove GP, Brown KK, et al. The idiopathic pulmonary fibrosis honeycomb cyst contains a mucocilary pseudostratified epithelium. PLoS One. 2013;8(3):e58658. Epub 2013/03/26. doi: 10.1371/journal.pone.0058658. PubMed PMID: 23527003; PubMed Central PMCID: PMCPMC3603941.

55. Helling BA, Gerber AN, Kadiyala V, Sasse SK, Pedersen BS, Sparks L, et al. Regulation of MUC5B Expression in Idiopathic Pulmonary Fibrosis. Am J Respir Cell Mol Biol. 2017;57(1):91–9. Epub 2017/03/09. doi: 10.1165/rcmb.2017-0046OC. PubMed PMID: 28272906; PubMed Central PMCID: PMCPMC5516283.

56. Arason AJ, Jonsdottir HR, Halldorsson S, Benediktsdottir BE, Bergthorsson JT, Ingthorsson S, et al. deltaNp63 has a role in maintaining epithelial integrity in airway epithelium. PLoS One. 2014;9(2):e88683. Epub 2014/02/18. doi: 10.1371/journal.pone.0088683. PubMed PMID: 24533135; PubMed Central PMCID: PMCPMC3922990.

57. Vanoni S, Tsai YT, Waddell A, Waggoner L, Klarquist J, Divanovic S, et al. Myeloid-derived NF-kappaB negative regulation of PU.1 and c/EBP-beta-driven pro-inflammatory cytokine production restrains LPS-induced shock. Innate Immun. 2017;23(2):175–87. Epub 2016/12/10. doi: 10.1177/1753425916681444. PubMed PMID: 27932520; PubMed Central PMCID: PMCPMC5563821.

58. Chinery R, Brockman JA, Dransfield DT, Coffey RJ. Antioxidant-induced nuclear translocation of CCAAT/enhancer-binding protein beta. A critical role for protein kinase A-mediated phosphorylation of Ser299. J Biol Chem. 1997;272(48):30356–61. Epub 1997/12/31. PubMed PMID: 9374525.

59. Coetzee SG, Coetzee GA, Hazelett DJ. motifbreakR: an R/Bioconductor package for predicting variant effects at transcription factor binding sites. Bioinformatics. 2015;31(23):3847–9. Epub 2015/08/15. doi: 10.1093/bioinformatics/btv470. PubMed PMID: 26272984; PubMed Central PMCID: PMCPMC4653394.

60. Yeh WC, Cao Z, Classon M, McKnight SL. Cascade regulation of terminal adipocyte differentiation by three members of the C/EBP family of leucine zipper proteins. Genes Dev. 1995;9(2):168–81. Epub 1995/01/15. PubMed PMID: 7531665.

61. Kono Y, Nishiuma T, Okada T, Kobayashi K, Funada Y, Kotani Y, et al. Sphingosine kinase 1 regulates mucin production via ERK phosphorylation. Pulm Pharmacol Ther. 2010;23(1):36–42. Epub 2009/10/20. doi: 10.1016/j.pupt.2009.10.005. PubMed PMID: 19835973.

62. Atherton HC, Jones G, Danahay H. IL-13-induced changes in the goblet cell density of human bronchial epithelial cell cultures: MAP kinase and phosphatidylinositol 3-kinase regulation. Am J Physiol Lung Cell Mol Physiol. 2003;285(3):L730–9. Epub 2003/06/10. doi: 10.1152/ajplung.00089.2003. PubMed PMID: 12794003.

63. Hewson CA, Edbrooke MR, Johnston SL. PMA induces the MUC5AC respiratory mucin in human bronchial epithelial cells, via PKC, EGF/TGF-alpha, Ras/Raf, MEK, ERK and Sp1-dependent mechanisms. J Mol Biol. 2004;344(3):683–95. Epub 2004/11/10. doi: 10.1016/j.jmb.2004.09.059. PubMed PMID: 15533438.

64. Zhu Y, Saunders MA, Yeh H, Deng WG, Wu KK. Dynamic regulation of cyclooxygenase-2 promoter activity by isoforms of CCAAT/enhancer-binding proteins. J Biol Chem. 2002;277(9):6923–8. Epub 2001/12/14. doi: 10.1074/jbc.M108075200. PubMed PMID: 11741938.

65. Pless O, Kowenz-Leutz E, Knoblich M, Lausen J, Beyermann M, Walsh MJ, et al. G9a-mediated lysine methylation alters the function of CCAAT/enhancer-binding protein-beta. J Biol Chem. 2008;283(39):26357–63. Epub 2008/07/24. doi: 10.1074/jbc.M802132200. PubMed PMID: 18647749; PubMed Central PMCID: PMCPMC3258912.

66. Roy SK, Hu J, Meng Q, Xia Y, Shapiro PS, Reddy SP, et al. MEKK1 plays a critical role in activating the transcription factor C/EBP-beta-dependent gene expression in response to IFN-gamma. Proc Natl Acad Sci U S A. 2002;99(12):7945–50. Epub 2002/06/06. doi: 10.1073/pnas.122075799. PubMed PMID: 12048245; PubMed Central PMCID: PMCPMC123000.

67. Kinoshita S, Akira S, Kishimoto T. A member of the C/EBP family, NF-IL6 beta, forms a heterodimer and transcriptionally synergizes with NF-IL6. Proc Natl Acad Sci U S A. 1992;89(4):1473–6. Epub 1992/02/15. PubMed PMID: 1741402; PubMed Central PMCID: PMCPMC48473.

68. Roy MG, Livraghi-Butrico A, Fletcher AA, McElwee MM, Evans SE, Boerner RM, et al. Muc5b is required for airway defence. Nature. 2014;505(7483):412–6. Epub 2013/12/10. doi: 10.1038/nature12807. PubMed PMID: 24317696; PubMed Central PMCID: PMCPMC4001806.

69. Cain DW, O’Koren EG, Kan MJ, Womble M, Sempowski GD, Hopper K, et al. Identification of a tissue-specific, C/EBPbeta-dependent pathway of differentiation for murine peritoneal macrophages. J Immunol. 2013;191(9):4665–75. Epub 2013/10/01. doi: 10.4049/jimmunol.1300581. PubMed PMID: 24078688; PubMed Central PMCID: PMCPMC3808250.

70. Tamura A, Hirai H, Yokota A, Sato A, Shoji T, Kashiwagi T, et al. Accelerated apoptosis of peripheral blood monocytes in Cebpb-deficient mice. Biochem Biophys Res Commun. 2015;464(2):654–8. Epub 2015/07/15. doi: 10.1016/j.bbrc.2015.07.045. PubMed PMID: 26168729.

71. Satoh T, Nakagawa K, Sugihara F, Kuwahara R, Ashihara M, Yamane F, et al. Identification of an atypical monocyte and committed progenitor involved in fibrosis. Nature. 2017;541(7635):96–101. Epub 2016/12/22. doi: 10.1038/nature20611. PubMed PMID: 28002407.

72. Silva MA, Bercik P. Macrophages are related to goblet cell hyperplasia and induce MUC5B but not MUC5AC in human bronchus epithelial cells. Lab Invest. 2012;92(6):937–48. Epub 2012/03/07. doi: 10.1038/labinvest.2012.15. PubMed PMID: 22391959.

73. Chen Y, Zhao YH, Di YP, Wu R. Characterization of human mucin 5B gene expression in airway epithelium and the genomic clone of the amino-terminal and 5’-flanking region. Am J Respir Cell Mol Biol. 2001;25(5):542–53. Epub 2001/11/20. doi: 10.1165/ajrcmb.25.5.4298. PubMed PMID: 11713095.

74. Viart V, Varilh J, Lopez E, Rene C, Claustres M, Taulan-Cadars M. Phosphorylated C/EBPbeta influences a complex network involving YY1 and USF2 in lung epithelial cells. PLoS One. 2013;8(4):e60211. Epub 2013/04/06. doi: 10.1371/journal.pone.0060211. PubMed PMID: 23560079; PubMed Central PMCID: PMCPMC3613372.

75. Vittal R, Mickler EA, Fisher AJ, Zhang C, Rothhaar K, Gu H, et al. Type V collagen induced tolerance suppresses collagen deposition, TGF-beta and associated transcripts in pulmonary fibrosis. PLoS One. 2013;8(10):e76451. Epub 2013/11/10. doi: 10.1371/journal.pone.0076451. PubMed PMID: 24204629; PubMed Central PMCID: PMCPMC3804565.

