## Supplementary files for "A MUC5B polymorphism associated with Idiopathic Pulmonary Fibrosis mediates overexpression through decreased CpG methylation and C/EBPβ transcriptional activation"

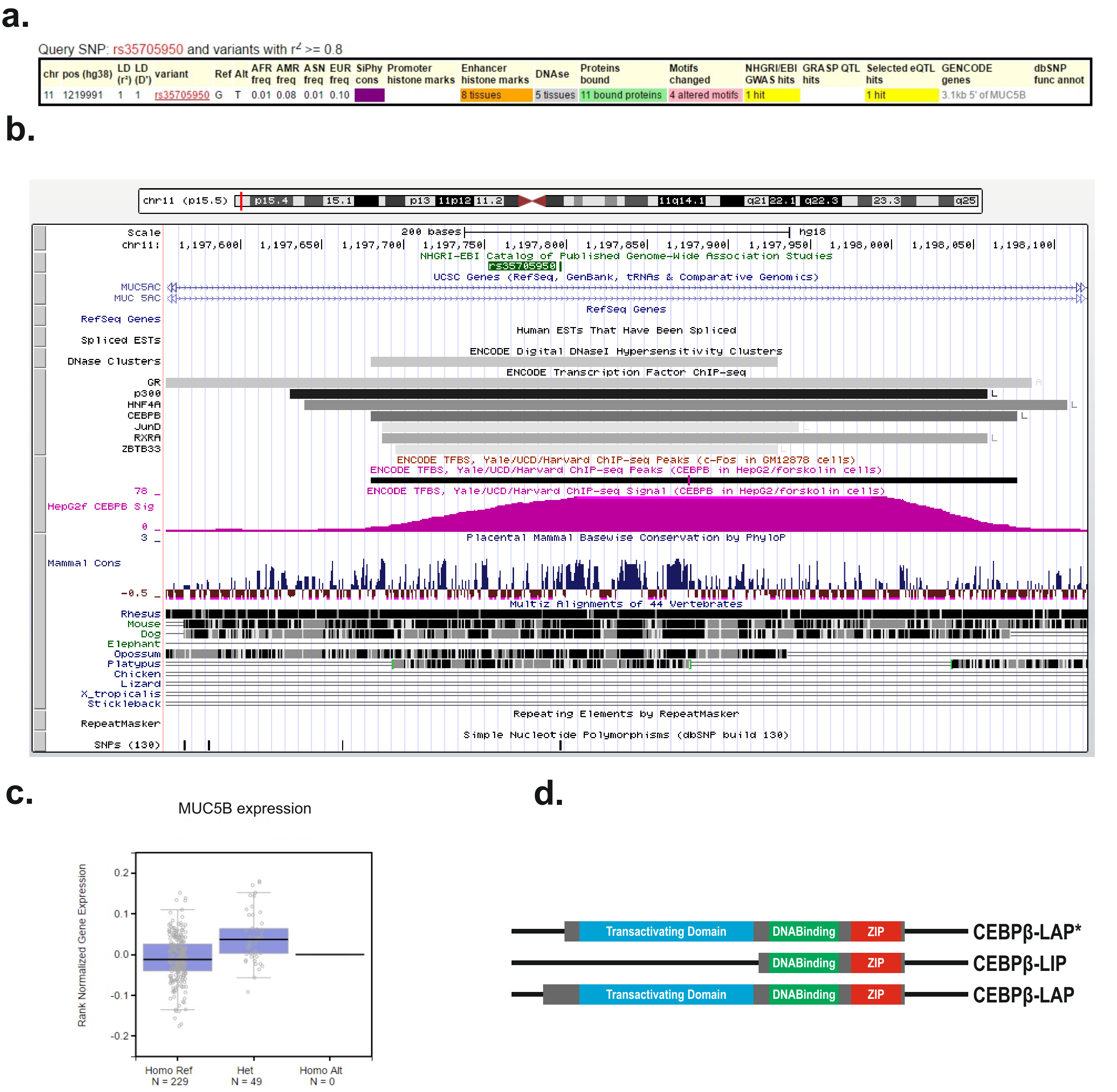

**S1 Fig. Schematic representation of rs35705950 on genome browser.** Descriptive data for rs35705950 polymorphism obtained from GTEx database (a) and UCSC genome browser (b). c) Differential *MUC5B* expression between homozygous [G/G] and heterozygous [G/T] carriers of rs35705950. Data obtained from GTEx database based on human genotyped samples. d) A schematic representation of the different C/EBPβ isoforms.

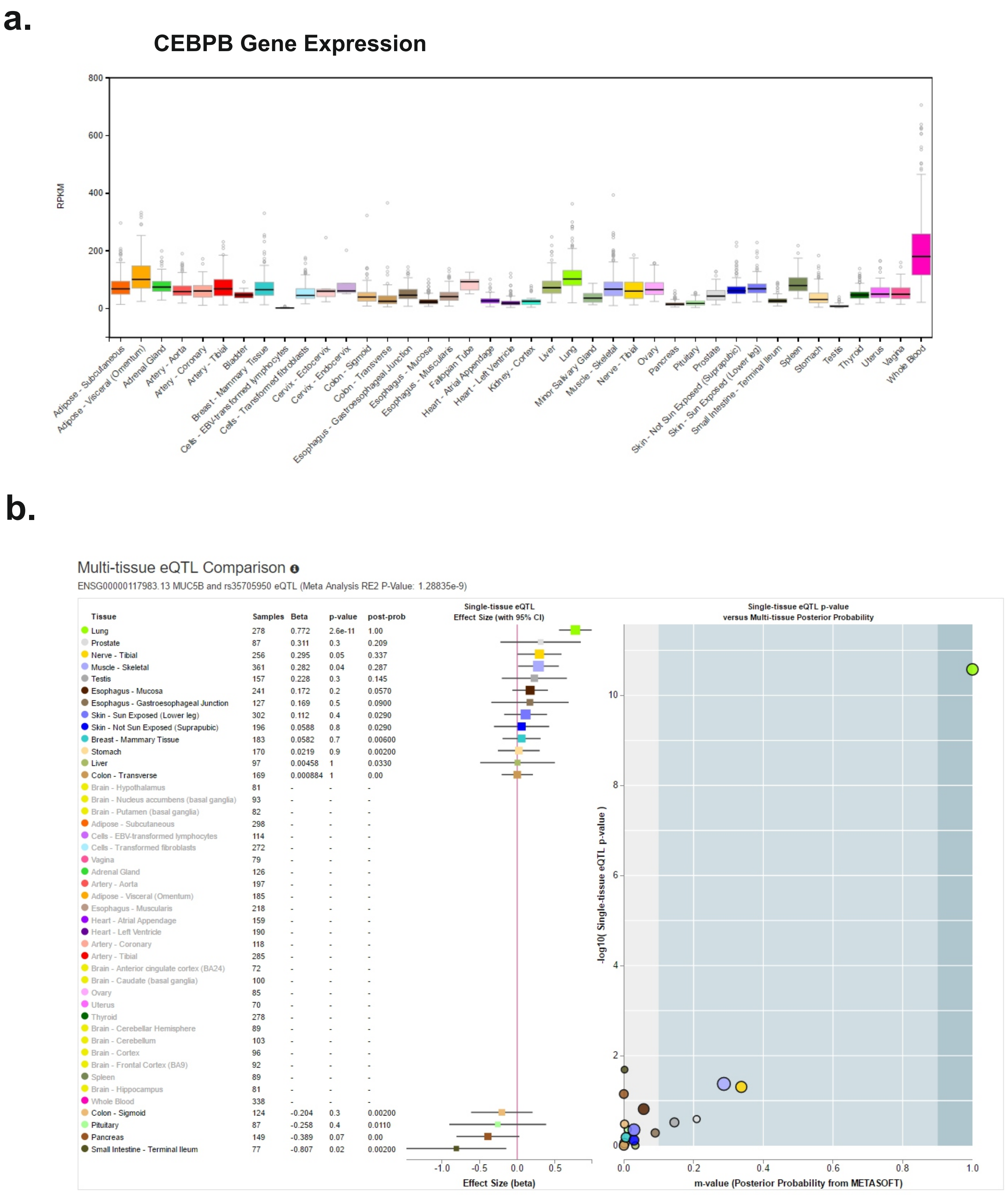

**S2 Fig. Analysis of CEBPB as a candidates to bind to the MUC5B *cis*-regulatory domain through the T allele.** a)C/EBPβ organ related expression from GTEx. b) rs35705950 polymorphism multi-tissue eQTL plot from GTEx.

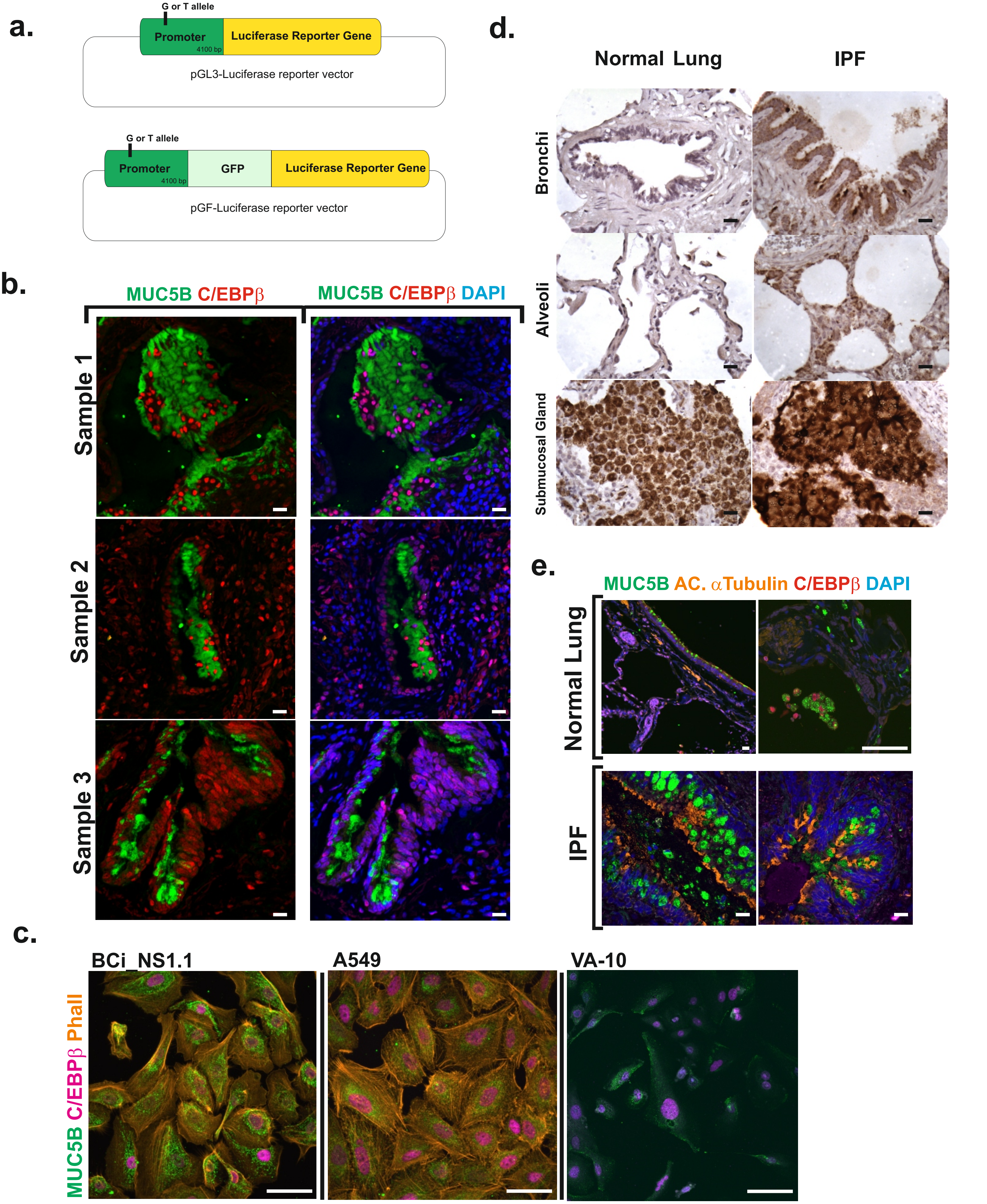

**S3 Fig. MUC5B and C/EBPβ are co-expressed in IPF and healthy lung samples.** a) Schematic view of Luciferase reporter vectors used in Figure 2b (top) and Figure 1a (bottom). b) C/EBPβ and MUC5B are co-expressed in IPF epithelial cells. The pictures are a representation of 3 different samples. c) C/EBPβ (red) and MUC5B (green) are co-expressed in Bci_NS1.1, A549 and VA-10 cell lines. Nucleus (blue) was stained with DAPI. d) MUC5B is overexpress in IPF compared to control samples. e) Comparison of MUC5B and C/EBPβ expression in IPF and control samples. Scale bars = 50µm.

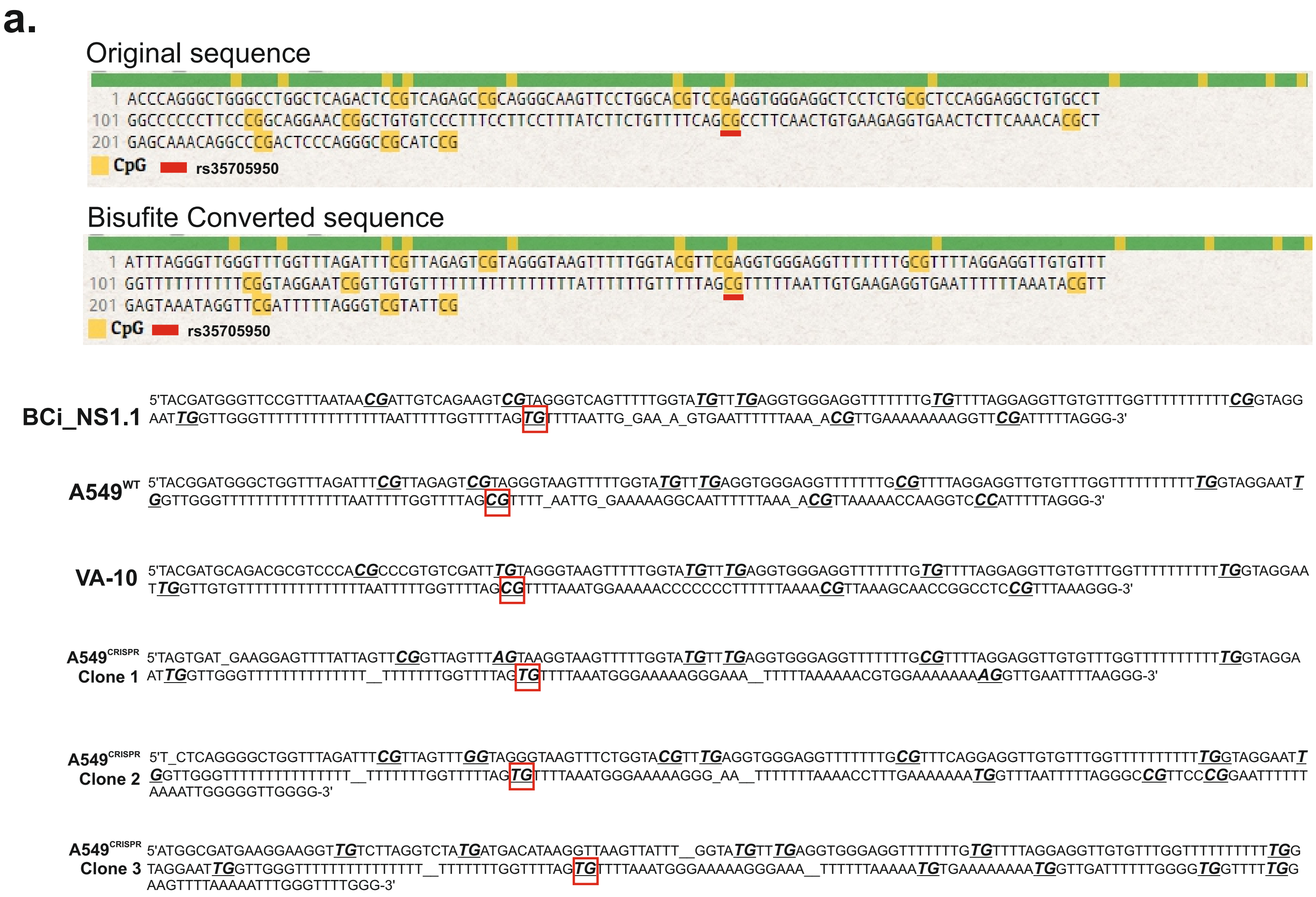

**S4 Fig. Sequencing data of bisulfite sequencing experiment.** a) Bisulfite sequencing of BCi_NS1.1, A549, VA10 and A549[CRISPR] clones (1-5). Yellow box represents CpG site and red line the rs35705950 position, overlapping a CpG site. Bisulfite expected converted sequence was obtained from ZymoResearch Bisulfite Online Tool.

a.

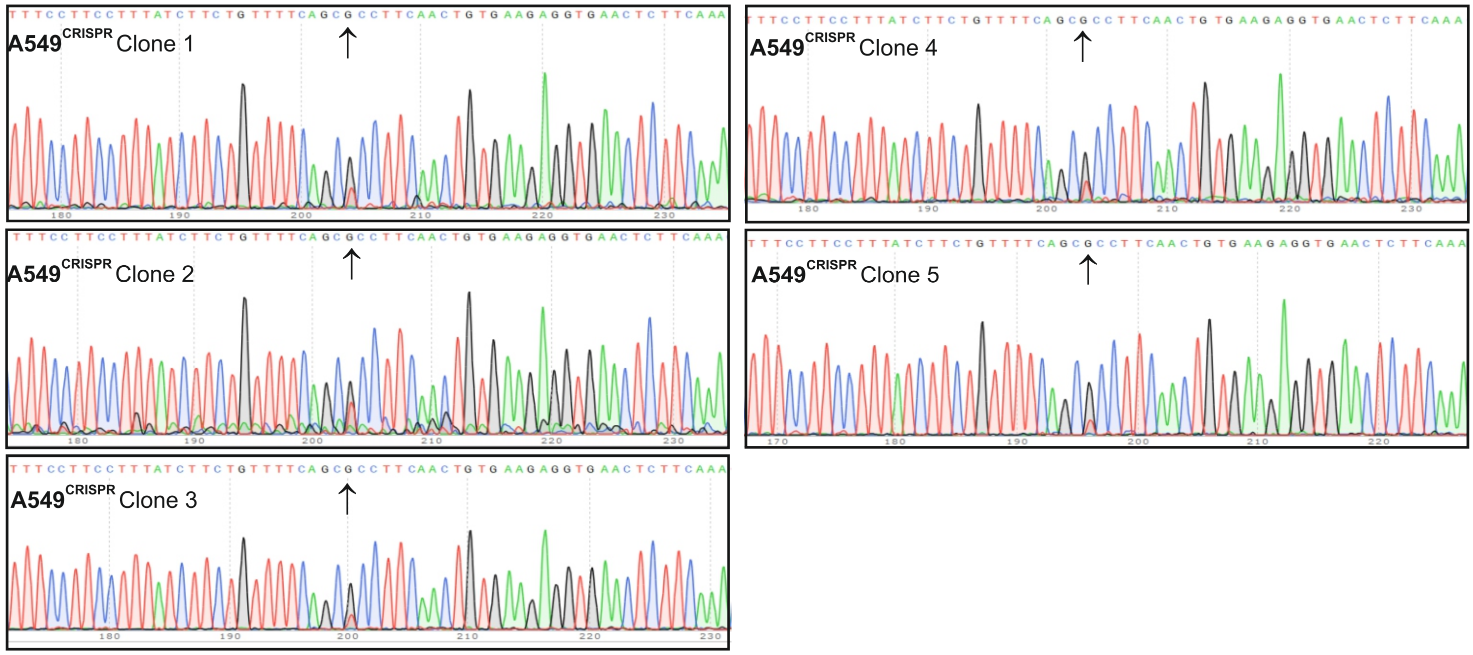

**S5 Fig. Sequencing data of CRISPR-Cas9 experiments.** a) Genotype of A549[CRISPR] clones.

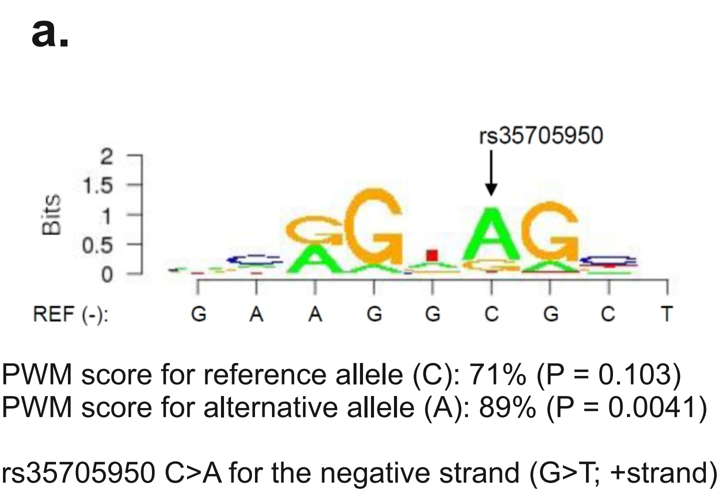

b.

CEBP; Hsapiens-cispb_1.02-M3036_1.02 (CIS-BP database)

| | 1 | 2 | 3 | 4 | 5 | 6 | 7 | 8 | 9 |
|---|---|---|---|---|---|---|---|---|---|
| A | 0,40 | 0,20 | 0,49 | 0,08 | 0,25 | 0,80 | 0,14 | 0,10 | 0,30 |
| C | 0,24 | 0,54 | 0,00 | 0,00 | 0,04 | 0,00 | 0,00 | 0,53 | 0,25 |
| G | 0,18 | 0,13 | 0,51 | 0,91 | 0,16 | 0,17 | 0,84 | 0,18 | 0,22 |
| T | 0,18 | 0,13 | 0,00 | 0,01 | 0,55 | 0,03 | 0,02 | 0,19 | 0,24 |

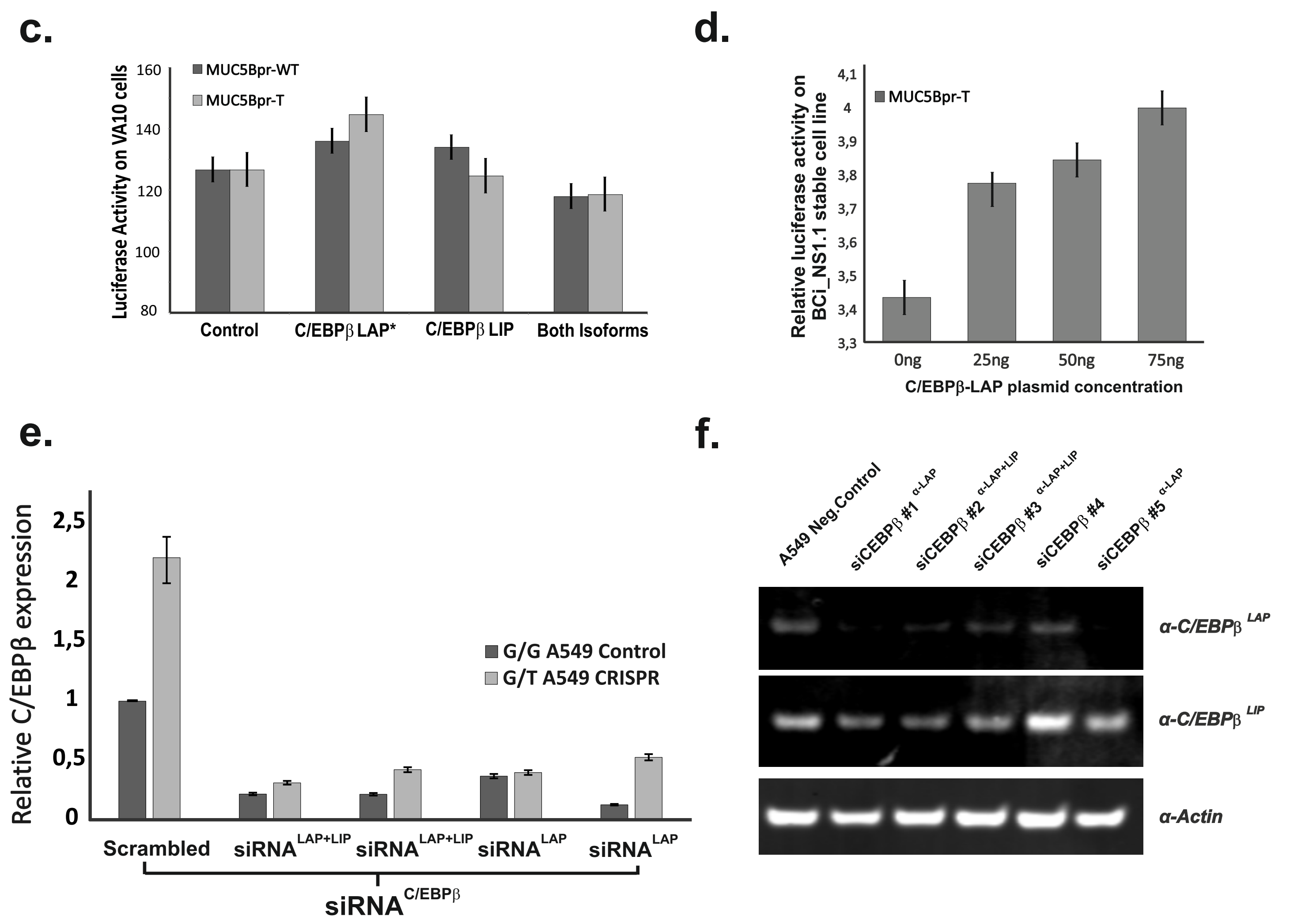

**S6 Fig. C/EBPβ increases differential MUC5B expression between WT and T alleles.** a) The alternative allele for rs35705950 contains a candidate binding site for C/EBPβ on the negative strand. C/EBPβ motif derived from MotifDb (Bioconductor package for R). REF(-)=reference sequence for the negative strand. The position weight matrix (PWM) score for reference allele is 71% while alternative allele has a PWM score of 89%. b) The alternative allele for rs35705950 contains the crucial nucleotides for the binding of C/EBPβ on the negative strand. Each number indicates the nucleotide position in the standard binding site motif. The importance of the nucleotide is indicated by a green scale. Dark green indicates critial nucleotide for CEBPB binding while yellow means the opposite. c) The three C/EBPβ isoforms were transfected individually into VA10 cell line with non-significant difference. d) Three different concentration of C/EBPβ-LAP plasmid were transfected in BCi_NS1.1 luciferase stable cell line, showing a dose dependent MUC5B expression through T allele. e) Corroboration of C/EBPβ relative expression after siRNA knock down in Figure 4c for A549 CRIPSR cell line and A549 Cas9-control (WT). f) Western blot shows C/EBPβ expression as a corroboration of C/EBPβ siRNA knock down in Figure 4d. siC/EBPβ#4 was discarded and not used on the experiments due to the low knock-down ratio.

### S1 Table. Summary of primer sequences.

| ChIP | Forward | Reverse |
|---|---|---|
| ChIP1-M5Bpr | AGCTATTGAGACATCCCGGA | GCTGTGTCCCTTTCCTTCCT |
| ChIP2-M5Bpr | GAGTCGGGCCTGTTTGCT | AACCGGCTGTGTCCCTTT |
| ChIP3-M5Bpr | TGGGAGTCGGGCCTGTTT | CAGGAACCGGCTGTGTCC |
| ChIP4-M5Bpr | CTGCAGATGACGCTGTCTGT | GGGGCCCCAGCTTATGTAG |
| Negative Control - GAPDH | TCG AAC AGG AGG AGC AGA GAG CGA | TAC TAG CGG TTT TAC GGG CG |
| Positive Control - TNFAIP3 | GCTGTTGCTCAATTGCTAGTC | CTTCTTGTGCTTACTTTCAGTTCTT |
| **CRISPR** | **Forward** | **Reverse** |
| Sequencing 1 | ATGCTACTGGAAGCCTCGAA | CATCAGCTCCCAGGCACT |
| Sequencing 2 | ATGCTACTGGAAGCCTCGAA | CATCAGCTCCCAGGCACT |
| Sequencing 3 | GTTGGACCACAGGCACTGA | CTGCACAGCGACGTGAAC |
| Sequencing 4 | CTGCAGATGACGCTGTCTGT | GGGGCCCCAGCTTATGTAG |
| Sequencing 5 | GAATTCATGCTACTGGAAGCCTCGAA | TCTAGAGGGTCTGTCCCCAGAAGAAC |
| MUC5B gRNA1 | AAAACTGGGAGTCGGGCCTGTTTGC | ACACCGCAAACAGGCCCGACTCCCAG |
| MUC5B gRNA2 | AAAACCAAACACGCTGAGCAAACAGC | ACACCGCTGTTTGCTCAGCGTGTTTG |
| MUC5B gRNA4 | ACACCGGGAGTCGGGCCTGTTTGCT | AAAACAGCAAACAGGCCCGACTCCC |
| MUC5B gRNA5 | AAAACGATGCGGCCCTGGGAGTCGGC | ACACCGCCGACTCCCAGGGCCGCATC |
| Homologous region 1 | TTATCTTCTGTTTTCAGCTCCTTCAACTGTGAAGAGGTGAACTCTTCAAACACGCTGAGCAAACAGGCCCGACTCCCA | |
| | GGGCCGCATCCGGGATGTCTCAATAGCTGTGGCCTTGACGTCCACCTCGGACCCCTGCCCCGGACCCAGCCCA | |
| Homologous region 2 | AGGTGGACGTCAAGGCCACAGCTATTGAGACATCCCGGATGCGGCCCTGGGAGTCGGGCCTGTTTGCTCAGCGTG | |
| | TTTGAAGAGTTCACCTCTTCACAGTTGAAGGAGCTGAAAACAGAAGATAAAGGAAGGAAAGGGACACAGCCGGTTC | |
| **Clonning primers** | **Forward** | **Reverse** |
| SEQ1 | ATGCTACTGGAAGCCTCGAA | CATCAGCTCCCAGGCACT |
| SEQ2 | GGCTCTGAGCAGACCAAGAG | CTCAGGCAGCTCCTCTGTC |
| SEQ3 | GTTGGACCACAGGCACTGA | CTGCACAGCGACGTGAAC |
| SEQ4 | AGCCATGAGGGGTGACAG | TGACGAGCGTCATCTACAGG |
| SEQ5 | CTGCAGATGACGCTGTCTGT | GGGGCCCCAGCTTATGTAG |
| MUC5Bpr-pGL3-MluI-NheI | ACGCGTATGCTACTGGAAGCCTCGAA | GCTAGCGGGTCTGTCCCCAGAAGAAC |
| MUC5Bpr-pGF-XbaI-EcoRI | GAATTCATGCTACTGGAAGCCTCGAA | TCTAGAGGGTCTGTCCCCAGAAGAAC |
| rs35705950_SEQ_587bp | CGGGTTCTGTGTGGTCTAGG | GCATCAGCGAGATAGCGTTT |
| Site Directed Mutagenesis [T] | CCTACGAAGGtGACAAAGAGTCCG | TGTGGCAGCGTCCTGGGA |
| Site Directed Mutagenesis [T] | CCTACGAAGGaGACAAAGAGTCCG | TGTGGCAGCGTCCTGGGA |
| **Bisulfite sequencing** | **Fordward** | **Reverse** |
| Bisulfite Sequencing | TTTGGTTAGAATGAGGGATAGTGAT | CAAAACCACAACTATTAAAACATCC |
| **qRT-PCR** | **IDT Identification number** | |
| MUC5B | Hs.PT.5822513172g | |
| MUC5AC | Hs.PT.5115096441g | |
| GAPDH | Hs.PT.39a.22214836 | |
| C/EBPβ | Hs.PT.5827185099g | |

#### S2 Table. Summary of antibodies.

| Protein | Identification number |
|---|---|
| MUC5B | ab87376 |
| MUC5AC | ab3649 |
| C/EBPβ | ab32358 |
| Acethylated α-tubulin | ab11323 |
| Phalloidin | a22283 |
| **ChIP** | **Identification number** |
| C/EBPβ | sc-150 |
| Rabbit (DA1E) mAb IgG XP® Isotype Control | #3900 |

#### S3 Table. Summary of plasmids.

| MUC5B promoter Cloning vectors | Catalogue Number | Company |
|---|---|---|
| pGL3-LUC reporter vector | E1751 | Promega |
| Renilla Reporter vector | E2231 | Promega |
| pGF-LUC-GFP Lentivector | TR010PA-N | System Bioscience |
| pGF-LUC-GFP Negative Control | TR000PA-1 | System Bioscience |
| psPAX2 | #12260 | Addgene |
| pMD2.G | #12259 | Addgene |
| **CRISPR Vectors** | **Catalogue Number** | **Company** |
| CAS9-Nickase | #51130 | Addgene |
| gRNA backbone (MLM3636) | #43860 | Addgene |
| **C/EBPβ overexpression** | **Catalogue Number** | **Company** |
| C/EBPβ LAP | #15738 | Addgene |
| C/EBPβ LIP | #15737 | Addgene |

### S4 Table. Allele-specific predictions for DNA binding protein affinity.

**A) DNA binding protein occupancy predictions for minor allele (sites created by minor allele)**

| Chr | Start_hg 38 | End_hg38 | Strand | Width | Gene | PWM motif name | DNA(minor allele) 5´>3´ | DNA(major allele) 5´>3´ | P-value (minor allele) | P-value (major allele) | PWM score (minor allele) | PWM score (major allele) | #critical nucleotides in sequence specificity | #critical nucleotides matched for minor allele | #critical nucleotides matched for major allele | providerID | GTRD ChIP-seq experimental support |
|---|---|---|---|---|---|---|---|---|---|---|---|---|---|---|---|---|---|
| chr11 | 1219987 | 1219996 | - | 10 | MYOD1 | Hsapiens-cispb_1.02-M3589_1.02 | GAAGGAGCTG | GAAGGCGCTG | 0,002347887 | 0,063471221 | 0,82949113 | 0,625627867 | 4 | 3 | 2 | MA0056823_1.02 | |
| chr11 | 1219987 | 1219994 | - | 8 | TFAP2B | Hsapiens-cispb_1.02-M5919_1.02 | AGGAGCTG | AGGCGCTG | 0,002896477 | 0,023817417 | 0,789397453 | 0,649224842 | 6 | 4 | 3 | MA0033086_1.02 | |
| chr11 | 1219987 | 1219994 | - | 8 | ATF1 | Hsapiens-cispb_1.02-M6152_1.02 | AGGAGCTG | AGGCGCTG | 0,003664218 | 0,03652004 | 0,811929183 | 0,661889531 | 5 | 3 | 2 | MA0110391_1.02 | HepG2 |
| chr11 | 1219987 | 1219994 | - | 8 | TFAP2B | Hsapiens-cispb_1.02-M5918_1.02 | AGGAGCTG | AGGCGCTG | 0,004205126 | 0,037898484 | 0,779493324 | 0,637838924 | 7 | 5 | 4 | MA0033085_1.02 | |
| chr11 | 1219988 | 1219995 | - | 8 | ATF3 | Hsapiens-cispb_1.02-M4683_1.02 | AAGGAGCT | AAGGCGCT | 0,004573913 | 0,027278261 | 0,767395079 | 0,640170076 | 7 | 5 | 4 | MA0110551_1.02 | A549, HepG2 |
| chr11 | 1219987 | 1219996 | - | 10 | USF1 | Hsapiens-cispb_1.02-M4514_1.02 | GAAGGAGCTG | GAAGGCGCTG | 0,006052332 | 0,156073637 | 0,861249786 | 0,609814605 | 3 | 3 | 2 | MA0057014_1.02 | A549 |
| chr11 | 1219988 | 1219996 | - | 9 | CEBPB | Hsapiens-cispb_1.02-M3036_1.02 | GAAGGAGCT | GAAGGCGCT | 0,006261033 | 0,110942025 | 0,880897425 | 0,703883729 | 3 | 3 | 2 | MA0110670_1.02 | A549, Ishikawa, MCF7, LS180, HepG2 |
| chr11 | 1219990 | 1219996 | - | 7 | MYC | Hsapiens-cispb_1.02-M4645_1.02 | GAAGGAG | GAAGGCG | 0,006991304 | 0,1624 | 0,869137742 | 0,587662412 | 2 | 2 | 1 | MA0056906_1.02 | MCF7, HCT-116 |
| chr11 | 1219985 | 1219995 | + | 11 | TFE3 | Hsapiens-cispb_1.02-M5930_1.02 | TTCAGCTCCTT | TTCAGCGCCTT | 0,008488772 | 0,059875068 | 0,72059545 | 0,574075057 | 6 | 4 | 3 | MA0056463_1.02 | HepG2 |
| chr11 | 1219987 | 1219993 | + | 7 | XBP1 | Hsapiens-cispb_1.02-M5953_1.02 | CAGCTCC | CAGCGCC | 0,00909075 | 0,137443074 | 0,771861852 | 0,543320634 | 4 | 3 | 2 | MA0110275_1.02 | |

**B) DNA binding protein occupancy predictions for major allele (sites disrupted by minor allele)**

| Chr | Start_hg 38 | End_hg38 | Strand | Width | Gene | PWM motif name | DNA(major allele) 5´>3´ | DNA(minor allele) 5´>3´ | P-value (major allele) | P-value (minor allele) | PWM score (major allele) | PWM score (minor allele) | #critical nucleotides in sequence specificity | #critical nucleotides matched for major allele | #critical nucleotides matched for minor allele | providerID | GTRD ChIP-seq experimental support |
|---|---|---|---|---|---|---|---|---|---|---|---|---|---|---|---|---|---|
| chr11 | 1219986 | 1219995 | - | 10 | BATF3 | Hsapiens-cispb_1.02-M5302_1.02 | AAGGCGCTGA | AAGGAGCTGA | 0,001052174 | 0,018313043 | 0,876051759 | 0,705091011 | 5 | 5 | 4 | MA0110407_1.02 | |
| chr11 | 1219987 | 1219996 | + | 10 | ATF7 | Hsapiens-cispb_1.02-M5293_1.02 | CAGCGCCTTC | CAGCTCCTTC | 0,001156522 | 0,024365217 | 0,88980748 | 0,692773956 | 4 | 4 | 3 | MA0110622_1.02 | |
| chr11 | 1219985 | 1219994 | - | 10 | ATF7 | Hsapiens-cispb_1.02-M5293_1.02 | AGGCGCTGAA | AGGAGCTGAA | 0,001643478 | 0,016434783 | 0,870390935 | 0,717593983 | 4 | 4 | 3 | MA0110622_1.02 | |
| chr11 | 1219986 | 1219995 | + | 10 | BATF3 | Hsapiens-cispb_1.02-M5302_1.02 | TCAGCGCCTT | TCAGCTCCTT | 0,001678261 | 0,02613913 | 0,853878554 | 0,682917805 | 5 | 4 | 3 | MA0110407_1.02 | |
| chr11 | 1219984 | 1219995 | - | 12 | MAFK | Hsapiens-cispb_1.02-M4626_1.02 | AAGGCGCTGAAA | AAGGAGCTGAAA | 0,001886957 | 0,03546087 | 0,833473668 | 0,679266958 | 6 | 5 | 4 | MA0110839_1.02 | |
| chr11 | 1219987 | 1219996 | - | 10 | ATF3 | Hsapiens-cispb_1.02-M4451_1.02 | GAAGGCGCTG | GAAGGAGCTG | 0,002487021 | 0,01354818 | 0,927145425 | 0,798705626 | 4 | 4 | 3 | MA0110547_1.02 | A549, HepG2 |
| chr11 | 1219987 | 1219995 | + | 9 | BHLHA15 | Hsapiens-cispb_1.02-M5304_1.02 | CAGCGCCTT | CAGCTCCTT | 0,002521739 | 0,040643478 | 0,89554907 | 0,700286069 | 3 | 3 | 2 | MA0057226_1.02 | |
| chr11 | 1219987 | 1219997 | - | 11 | BACH1 | Hsapiens-cispb_1.02-M2979_1.02 | TGAAGGCGCTG | TGAAGGAGCTG | 0,002895728 | 0,06024505 | 0,86703472 | 0,711142014 | 5 | 3 | 2 | MA0110520_1.02 | |
| chr11 | 1219988 | 1219995 | + | 8 | JUN | Hsapiens-cispb_1.02-M4531_1.02 | AGCGCCTT | AGCTCCTT | 0,002991382 | 0,03467047 | 0,857916001 | 0,726468184 | 5 | 3 | 2 | MA0110734_1.02 | A549 |
| chr11 | 1219987 | 1219995 | - | 9 | CREB1 | Hsapiens-cispb_1.02-M4012_1.02 | AAGGCGCTG | AAGGAGCTG | 0,00390445 | 0,012878597 | 0,852506546 | 0,778497331 | 5 | 4 | 3 | MA0110370_1.02 | HepG2, A549 |
| chr11 | 1219986 | 1219995 | + | 10 | ATF4 | Hsapiens-cispb_1.02-M5292_1.02 | TCAGCGCCTT | TCAGCTCCTT | 0,004217391 | 0,050930435 | 0,896391722 | 0,713469084 | 3 | 3 | 2 | MA0110433_1.02 | |
| chr11 | 1219987 | 1219996 | - | 10 | ARNTL | Hsapiens-cispb_1.02-M5290_1.02 | GAAGGCGCTG | GAAGGAGCTG | 0,004313043 | 0,0558 | 0,875387735 | 0,674579136 | 3 | 3 | 2 | MA0056860_1.02 | |
| chr11 | 1219986 | 1219994 | - | 9 | BHLHA15 | Hsapiens-cispb_1.02-M5304_1.02 | AGGCGCTGA | AGGAGCTGA | 0,004504348 | 0,064486957 | 0,864419653 | 0,668846291 | 3 | 3 | 2 | MA0057226_1.02 | |
| chr11 | 1219985 | 1219994 | + | 10 | ARNTL | Hsapiens-cispb_1.02-M5290_1.02 | TTCAGCGCCT | TTCAGCTCCT | 0,004686957 | 0,060756522 | 0,870657705 | 0,669849107 | 3 | 3 | 2 | MA0056860_1.02 | |
| chr11 | 1219986 | 1219994 | - | 9 | TFAP2A | Hsapiens-cispb_1.02-M5914_1.02 | AGGCGCTGA | AGGAGCTGA | 0,004946694 | 0,045322887 | 0,857369815 | 0,749877969 | 3 | 3 | 2 | MA0033197_1.02 | SK-BR-3, BT-474, MCF7, MDA-MB-453 |
| chr11 | 1219986 | 1219995 | - | 10 | ATF4 | Hsapiens-cispb_1.02-M5292_1.02 | AAGGCGCTGA | AAGGAGCTGA | 0,005730435 | 0,073478261 | 0,869578458 | 0,686655819 | 3 | 3 | 2 | MA0110433_1.02 | |
| chr11 | 1219988 | 1219996 | - | 9 | NHLH1 | Hsapiens-cispb_1.02-M3376_1.02 | GAAGGCGCT | GAAGGAGCT | 0,006278425 | 0,063584267 | 0,875485902 | 0,666122283 | 4 | 3 | 2 | MA0057116_1.02 | |
| chr11 | 1219990 | 1219997 | - | 8 | USF1 | Hsapiens-cispb_1.02-M4429_1.02 | TGAAGGCG | TGAAGGAG | 0,00673061 | 0,066375645 | 0,806119616 | 0,639436975 | 6 | 4 | 3 | MA0057011_1.02 | A549 |

**S3 Table. Allele-specific predictions for DNA binding protein affinity overlapping rs35705950.** a) Binding sites created by minor allele b) Binding sites disrupted by minor allele.
